# Enabling spatiotemporal regulation within biomaterials using DNA reaction-diffusion waveguides

**DOI:** 10.1101/2022.02.26.482105

**Authors:** Phillip J. Dorsey, Dominic Scalise, Rebecca Schulman

## Abstract

In multicellular organisms, cells and tissues coordinate biochemical signal propagation across length scales spanning microns to meters. Endowing synthetic materials with similar capacities for coordinated signal propagation could allow these systems to adaptively regulate themselves across space and over time. Here we combine ideas from cell signaling and electronic circuitry to design a biochemical waveguide that transmits information in the form of a concentration of a DNA species on a directed path. The waveguide can be seamlessly integrated into a soft material because there is virtually no difference between the chemical or physical properties of the waveguide and the material it is embedded within. We propose the design of DNA strand displacement reactions to construct the system and, using reaction-diffusion models, identify kinetic and diffusive parameters that enable super-diffusive transport of DNA species via autocatalysis. Finally, to support experimental waveguide implementation, we show how a sink reaction could mitigate the spurious amplification of an autocatalyst within the waveguide, allowing for controlled waveguide triggering. Chemical waveguides could facilitate the design of synthetic biomaterials with distributed sensing machinery integrated throughout their structure and enable coordinated self-regulating programs triggered by changing environmental conditions.

## 1. Introduction

The biomolecular components of cells are powerful computational tools. They serve as exquisite detectors of signaling molecules^1^, pathogens^2^ and metabolites^3,4^, orchestrate multistep chemical synthesis and catalysis, and self-assemble nanostructures^5^ or materials with unique structural properties^6^ and geometry^7^. Synthetic biomolecular sensors can detect concentrations of drugs in the blood in real time^8,9^, approaching the sensitivity with which cells detect substances. Similarly, engineered enzyme cascades can orchestrate multistage chemical reactions^10^, biomolecular assemblies can template electronic devices^11,12^, and therapeutics can sense local conditions and dispense medication at the right time^13–15^. Recently, engineers and nanotechnologists have sought to design synthetic materials capable of sensing, integrating, and transmitting spatial information in processes similar to the functions performed by vascularized tissues. Microfabricated systems composed of fluidic or pneumatic vasculature have been designed to coordinate and direct delivery of fuel or nutrients to various locations in soft polymer substrates^16^ in order to control actuation and growth and migration of cells in tissue scaffolds^17^. However, fluidic control mechanisms present several challenges for designing triggerable sensing, communication, and computation in material systems. Such systems often require tethers to external power sources or fuel depots that are difficult to integrate within the structure of the material.

Here we use molecular programming concepts from synthetic biology and DNA nanotechnology to leverage the dynamics of non-linear reaction networks coupled to diffusive transport of molecular species to show how to achieve super-diffusive transport of chemical signals. Reaction-diffusion waveguides or wires consist of a region within a hydrogel substrate that acts as an excitable medium, where an autocatalytic reaction propagates spatially in the form of a traveling wavefront. Multiple wires could be integrated within a substrate and insulated from one another using competitive reactions that restrict autocatalytic reactions to the specific paths of the different wires. Chemical reactions generally take seconds to hours to reach completion and can require minimally nano-to micro-liter volumes to ensure deterministic behavior. As such, chemical wires are not intended to compete with electronic wires in terms of speed or reliability. Instead, the wires we develop, biochemical waveguides, are designed to enable the coordination of biomolecular sensors, polymer actuators, biomaterials, and soft robots in millimeter-scale soft materials without the need for electronics.

The idea of controlling information propagation through chemical reactions and directing information processing stems from Alan Turing’s seminal research regarding the origins of pattern formation during morphogenesis; this work described how periodic spatial patterns of chemical species could arise from transient fluctuations within an initially homogenous system.^18^ Experimental implementations of the Belousov-Zhabotinsky reaction-diffusion system (specifically aerosol OT microemulsion and chlorite-iodide-malonic acid reaction systems) have been demonstrated as mechanisms for propagating chemical species across space.^19,20^ Recently, the growing research field of DNA nanotechnology has provided new routes of material programming, enabling the design of experimental oscillators and amplifiers composed of biomolecular components capable of interfacing with biological systems.

Synthetic biomolecular devices have been developed that can release or respond to nucleic acid (DNA or RNA) signals of 20-100 bases in length. These signals can start or stop molecular machines^21^ or catalysis^22^, and direct hydrogel^23,24^ or nanostructure^25,26^ self-assembly. Nucleic acid signals can also be released by aptamer or antibody sensors^27,28^. Molecular circuits can represent information as concentrations of nucleic acids in solution. Just as electronic circuits can, molecular circuits can perform complex computation by emulating the functions of Boolean logic gates to conduct mathematical operations^29^ and can implement neural networks for pattern recognition^30^. These circuits can classify or execute logic operations on multiple nucleic acid signal inputs, act as memory latches, or direct oscillatory cycles of signal activity^29,31,32^. Enzymatic mechanisms may also be coupled with nucleic acids to initiate and regulate spatiotemporal changes in a system. For example, Bauer et al. demonstrated the formation of traveling waves of RNA in vitro generated from the Qβ viral enzyme and a self-replicating RNA species in a liquid filled capillary.^33^ Zadorin et al. used a polymerase-exonuclease-nickase (PEN) enzyme circuit with a template DNA duplex to produce a traveling wavefront within a buffer-filled polystyrene channel.^34^ Similarly, Zambrano et al. implemented an enzymatic Predator-Prey reaction network within a microfluidic network to compute the shortest distance within the network from entrance to exit^35^. Here we explore the design of non-enzymatic chemical reaction networks for propagating molecular signals. Temperature-dependent and batch-dependent variation in catalytic activity is often observed with enzymatic systems, which can yield varying stimuli-responsiveness between otherwise identical architectures^36,37^. These challenges could thus be avoided by using enzyme-free chemical reaction networks, which facilitate scalable and predictable design of molecular architectures where such sensitivities are unacceptable, and aid construction of more complex DNA-programable systems capable of executing spatial computation.

To this end, Chaterjee et al. implemented surface immobilized DNA complexes to catalytically exchange signals via strand displacement between DNA dominoes over nanometer length scales.^38^ Joesaar et al. developed a Sender-Receiver network consisting of DNA-encapsulated proteinosomes organized into regularly spaced arrays within a microfluidic trapping device as a means of exchanging chemical signals spatially between specific locations via diffusing DNA species.^39^ These microcapsules possessed DNA-permeable BSA-NH_2_/PNIPAAM crosslinked membranes and contained non-enzymatic catalytic DNA strand displacement circuits; when triggered by an externally supplied DNA signal, the microcapsule array executed distributed logic operations by exchanging diffusing DNA signals between Sender and Receiver microcapsules. Yang et al. built upon this work by incorporating a photo-triggerable DNA strand displacement latch within Sender microcapsules to enable distributed logical operations triggered through spatial activation with a laser.^40^ These studies are excellent models for how synthetic DNA-based systems might enable distributed communication spatially. However, the design of triggerable non-enzymatic DNA reaction-diffusion systems capable of transmitting chemical signals spatially as part of an integrated biomaterial substrate and the identification of experimentally attainable kinetic and diffusive parameters for such systems to achieve a rate of signaling faster than that provided by diffusion alone remains understudied.

Synthetic biologists and nanotechnologists have also sought to understand how non-linear chemical behaviors might be adapted, using DNA, to program excitatory responses in systems for signal amplification and information exchange in well-mixed conditions. Such non-linear dynamics, often consisting of chemical oscillators or amplifiers, commonly incorporate autocatalysis as a mechanism of amplification^41–43^. Here, we extend these analyses to characterize how such systems could function to enable directed spatial propagation of a DNA input as a reaction-diffusion system, and more specifically, how a strand displacement amplification network that we adapted from Zhang et al.^44^ could achieve this functionality. Gines et al., Lysne et al., and Srinivas et al. previously investigated thresholded autocatalytic DNA-based reaction networks in well-mixed systems^41–43^. Gines et al. implemented an enzymatic sink reaction in which an exonuclease quenched spurious amplification during microRNA detection.^41^ Srinivas et al. designed a strand displacement thresholding reaction that consumed autocatalyst species produced by parasitic reactions in an oscillatory strand-displacement reaction network.^43^ Lysne et al. implemented a leak-mitigation strategy by designing small interfering domains that hybridized to sections of a Fuel oligonucleotide to sterically hinder a leak reaction occurring in the well-mixed Zhang autocatalytic strand displacement amplifier.^42^

In this work, we develop a novel means of biomolecular signal propagation within soft materials, an insulated reaction-diffusion waveguide that could allow for super-diffusive transport over distances of hundreds of microns to millimeters. The waveguide insulation prevents propagated signals from spreading beyond the waveguide and acts as a chemical sink. By simultaneously directing and restricting transport of a chemical species, this design could accommodate integration of multiple waveguides to transmit multiple signals in parallel or propagate several different molecular inputs within a single substrate without triggering crosstalk. We first determine what rates of spatial propagation could be achieved by autocatalytic waves using an abstract autocatalytic chemical reaction network composed of a series of coupled bimolecular reactions within a reaction-diffusion medium. We then design a thresholding and amplification quenching strategy to enable insulation of waveguides and simultaneously prevent spurious activation by undesired leak reactions between different DNA species, similar to the damping strategy originally postulated by Zhang and Seelig^45^ and the thresholding reaction implemented by Srinivas et al.^39^. Finally, we implement the abstract enzyme-free autocatalysis reaction network and thresholding reaction as a DNA strand displacement reaction network designed for use in spatial signal propagation and show by measuring its rates in experiments, that this reaction scheme could be used to operate a triggerable autocatalytic waveguide.

## 2. Materials & Methods

### 2.1 Reaction-diffusion waveguide model

We constructed models of the reaction-diffusion waveguide using Comsol Multiphysics finite element analysis software. Specific details regarding the models’ geometry, initial conditions, reactions, and diffusion coefficients are provided in the supporting information (S3) and Results section.

### 2.2 DNA strand displacement reactions

DNA strands were purchased from Integrated DNA Technologies (Coralville, IA). The annealing and purification protocol for double-stranded DNA complexes as well as the protocol for reaction progress monitoring using quantitative PCR are provided in supplementary materials, S1. Curve fitting analysis to determine reaction rate constants is discussed in supplementary materials, S4.

## 3 Results

### 3.1 Autocatalytic amplification enables super-diffusive transport of biochemical species in a reaction-diffusion waveguide model

A reaction-diffusion waveguide consisted of a path within a hydrogel where specific DNA molecules were conjugated to the hydrogel’s polymer network. We modeled waveguide behavior using a 2-dimensional simulation where the waveguide path consisted of a 2-dimensional area (**Figure 2a**). The molecules along the path were the reactants and fuel consumed in an autocatalytic reaction. Another region of the hydrogel in which a different set of molecules was conjugated, termed as the insulation, lined the edges of the waveguide. The insulation contained a high concentration of a DNA species conjugated to the network that acted to prevent the wave propagating along the waveguide from extending beyond its boundaries. The following abstract reactions were propagated along the waveguide:

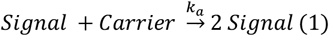

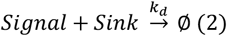

The Signal species served as a trigger for the autocatalytic reaction cascade and the operation of the waveguide involved the spatial propagation of a high concentration of Signal. Our model included a site for triggering wave propagation of Signal through the waveguide (location A in **Figure 1a**). When Signal was present at location A it could diffuse into the core wire domain and react with Carrier in the waveguide’s path to produce more Signal. The model of this process was designed to predict experiments that would be used to characterize a waveguide’s function. In such experiments, location A would contain Signal that was covalently linked to the hydrogel and thus could not diffuse into the waveguide. Shining UV light or another spatially modulated stimulus on location A would cleave the bonds connecting Signal to location A, allowing Signal to diffuse and trigger the waveguide’s operation. The waveguide propagates a chemical wave when Signal is added, because Signal can react with Carrier autocatalytically to produce 2 molecules of Signal (Reaction 1). This Signal can then diffuse and react with more Carrier, and thus generate more Signal. To make it possible to remove Signal from the waveguide after it was produced, we added Reaction 2, in which a Sink molecule reacts rapidly with Signal to convert it into waste. Carrier and Sink were immobilized within the waveguide core.

**Figure 1.**
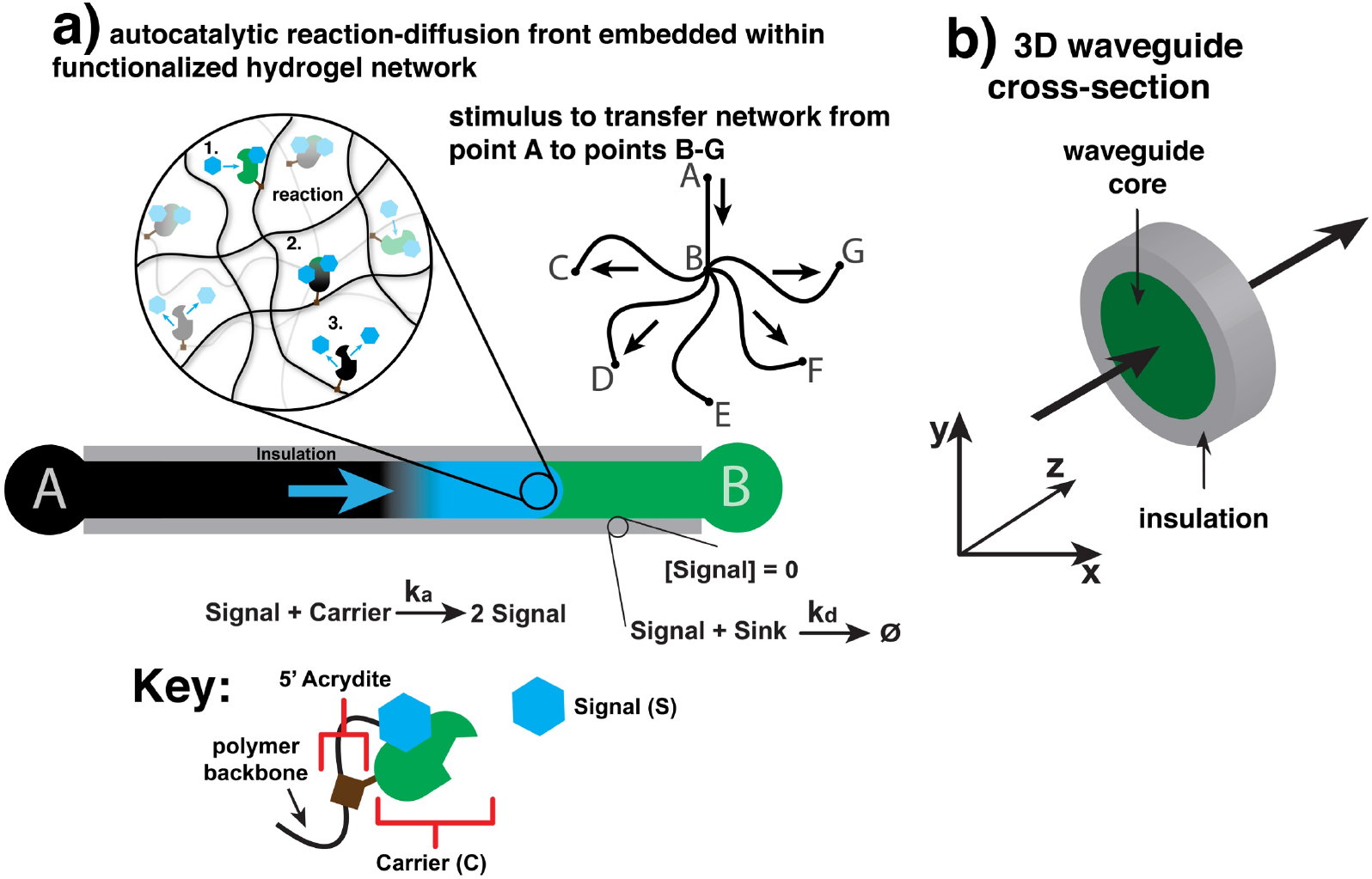
Schematic: Design and function of a reaction-diffusion waveguide in a hydrogel. a) A chemical wave of Signal is propagated between points A and B via an autocatalytic reaction that makes copies of Signal from a Carrier species that is crosslinked to the hydrogel network. Such a system could be used to route chemical signals simultaneously between multiple points in space: 1) Signal reacts with patterned Carrier, 2) Carrier transitions into its release state, 3) Carrier releases 2 Signal molecules. b) Schematic cross section of a 3-dimensional waveguide showing its core where autocatalysis occurs and the insulation surrounding it which prevents Signal from diffusing from the waveguide.

**Figure 2.**
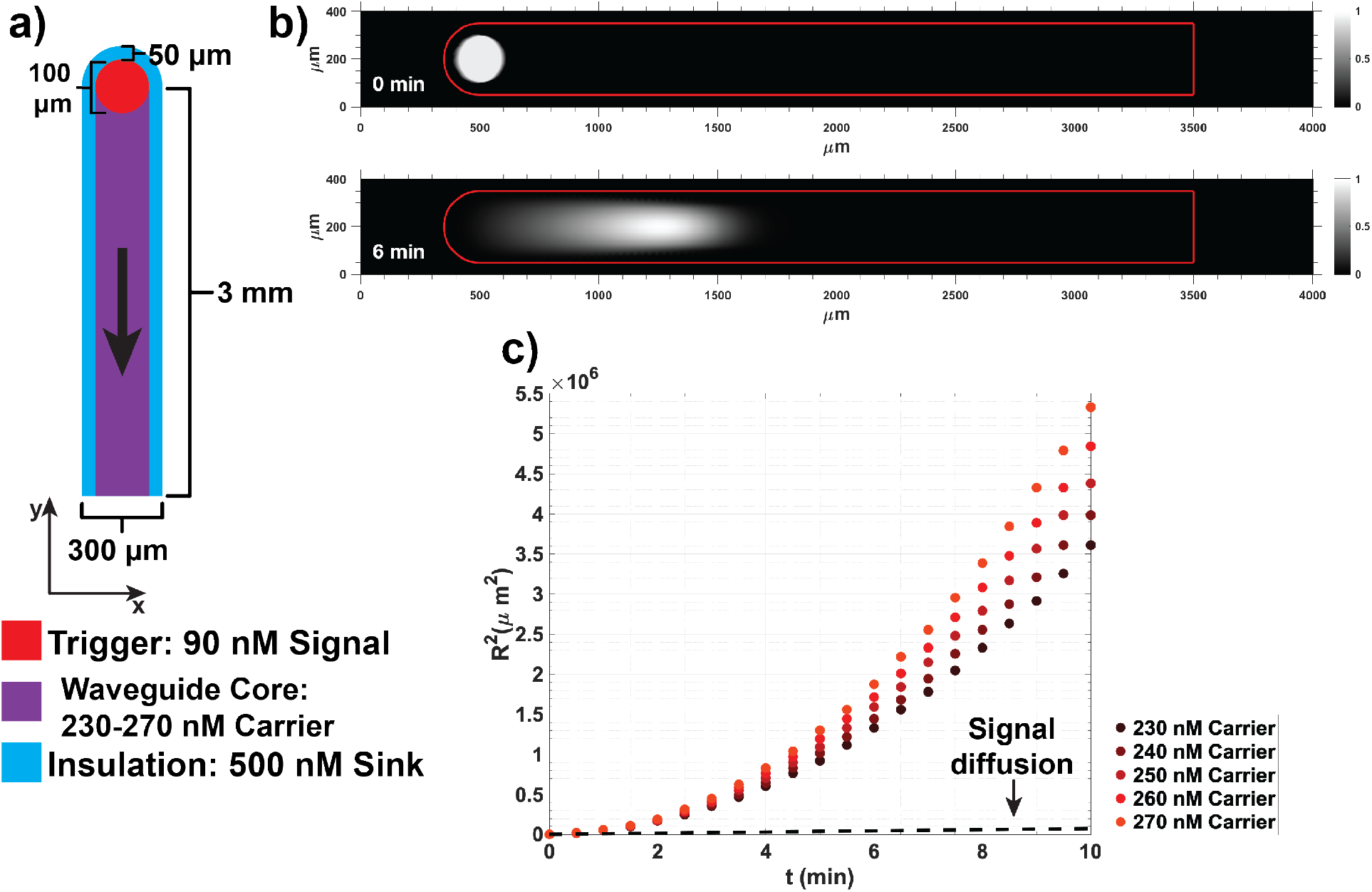
An idealized model of a reaction-diffusion waveguide. a) Geometry of the waveguide model and initial concentrations used for the amplification reaction. b) Top, the initial distribution of Signal before triggering and bottom, the distribution of Signal after 6 minutes of wave-front propagation down the waveguide. Both surface plots show concentrations normalized to maximum Signal concentration; the red frame indicates the boundaries of the waveguide. c) Squared wavefront displacement vs. time across 5 different Carrier concentrations. The dashed black line shows the squared displacement of a 42 nucleotide-sized single stranded DNA molecule over time resulting from diffusion alone.

The simplest method of transmitting information in the form of a particular molecular species between two points in space is to allow molecules to passively diffuse from a region of higher concentration to one of lower concentration. The characteristic time for this process to occur over a distance L scales with *O(L*^*2*^) according to Fick’s 2^nd^ law. Coupling a reaction to this diffusive process could decrease the time over which information is transmitted. Specifically, if a diffusing molecule, Signal, reacts with a patterned path of Carrier molecules to create copies of itself (as in Reaction 1), then Signal will form a moving wave in which Signal diffuses and also amplifies itself. With certain rates of reaction and diffusion, this process can produce a stable traveling wave. Such a wave forms when the autocatalytic reactions change the shape and magnitude of the Laplacian at the leading edge of the wavefront in such a way as to yield a linear rate of displacement with respect to time (assuming a constant concentration of Carrier along the waveguide path). The existence of a stable asymptotic traveling wavefront and the analytical relationship between reaction rate, diffusivity and wave velocity can be elucidated by adapting the Fisher-Kolomogorov-Petrovsky-Piskunov (FKPP) treatment^46–48^ of a one-dimensional reaction-diffusion process for the autocatalytic network described above. The full derivation of the equation is provided in section S5 of the supporting information. The final expression for the minimum rate of displacement for the autocatalytic front in the presence of Sink is:

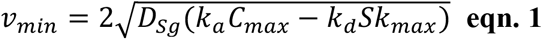

where *D*_*sg*_ is the diffusion coefficient of the Signal, *C*_*max*_ and *Sk*_*max*_ is the initial concentration of Carrier and Sink patterned within the core waveguide path, and *k*_*a*_ and *k*_*d*_ are the bimolecular rate constants for the abstract amplification and sink reactions respectively.

A key result of the FKPP analysis of the abstract amplification scheme is that the square of the effective change in displacement of the autocatalyst species in one-dimensional space (*δL*^2^) over time (*δt*) is proportional to the square of the change in time, *δL*^2^ ∝ *δt*^2^ * 4*D*_*Sg*_; this results in a power law dependence between *δL*^2^ and *δt* and a super-diffusive regime of Signal transport where the exponent of *δt* is greater than 1. For diffusion in the absence of any reaction, *δL*^2^ ∝ 2*D*_*A*_*δt*, which yields a linear relationship between the square of the displacement and time.

To verify that this idealized reaction-diffusion amplifier could achieve super-diffusive transport of an autocatalyst species, we simulated its behavior in a reaction-diffusion model. The model was implemented using Comsol Multiphysics (Transport of Dilute Species physics node). The waveguide was 3000 µm long and 300 µm wide (**Figure 2a**). The insulation surrounding the edges of the waveguide was 50 µm wide. In the model, the domain holding the initial Signal stimulus to trigger the system at one end of the waveguide (location A in Figure 1) consisted of a 100 µm radius circle.

Our first analysis modeled abstract reactions 1 and 2 and considered wave propagation when Sink was patterned within the insulation but not within the core waveguide path where Carrier was localized. The primary goal of these simulations was to determine whether idealized reactions 1 and 2 could form a stable traveling wave using net reaction rates that could be experimentally implemented for a toehold-mediated DNA strand displacement reaction network. A secondary goal was determine the form and speed of a traveling wave using an experimentally measured diffusion coefficient for short DNA molecules in poly(ethylene glycol) diacrylate hydrogels in order to show that these biomaterials could be used as substrate for the reaction network. The bimolecular rate constants for the modeled reactions were selected using measured rate constants for toehold mediated strand displacement reactions at 25º C in standard buffer conditions^49^. Strand displacement toeholds typically range in size from 0 to 7 nucleotides. Above toeholds of 7 nucleotides, the magnitude of the bimolecular rate constant saturates. Therefore, in order to design an amplifier that reacted at the fastest rate possible, we designed these reactions to occur with rate constants at the upper end of this scale. Specifically, we chose k_a_ to be 2×10^5^ M^-1^ s^-1^, which is on the same order of magnitude as the rate constant for a 6-nucleotide (nt) toehold strand displacement reaction; k_d_ was 4×10^6^ M^-1^ s^-1^, which is on the same order of magnitude as the rate constant for a standard 7-nt toehold reaction^49^. To ensure that the Sink reaction could successfully perform its function of restricting amplification to the waveguide, we set the rate constant for its reaction with Signal to be an order of magnitude higher than the rate constant for the Carrier and Signal reaction. Signal was assigned a diffusion coefficient of 60 µm^2^ sec^-1^, a typical value for the diffusivity of a 42-nt single stranded (ss) DNA oligonucleotide in poly(ethylene glycol) diacrylate (M_n_ = 575) hydrogels^50^. Sink and Carrier were immobilized within the waveguide insulation and waveguide core respectively (*i*.*e*. their diffusion constants were 0). The concentration of Sink in the insulation was 500 nM. The initial concentration of Signal within the triggered domain was 90 nM. The initial concentration of Carrier in the waveguide varied between 230 nM to 270 nM by steps of 10 nM in each simulation.

Across all Carrier conditions tested, we observed the formation of a stable Signal wavefront that traveled along the waveguide as it consumed Carrier (**Figure 2b and c**). Additionally, the wave was constrained to the waveguide; it did not spread beyond the insulation. We calculated the displacement of the wavefront in the center axis of the waveguide over time (**Figure 2c**) and observed that the square of the displacement, R^2^, was proportional to t^α^, with α > 1, indicating that the idealized system had achieved super-diffusive transport of Signal. The dashed black line in **Figure 2c** indicates the square of the displacement resulting from simple diffusion of a DNA oligonucleotide in one dimension.

While R^2^ for the reaction network grows exponentially with time, R^2^ in the case of simple diffusion grows linearly with time. Plots of R^2^ vs. time on a logarithmic plot yielded straight lines across all Carrier concentrations (**Figure 3**), where the slope of the line was α. Across all Carrier concentrations, the average value of α calculated from the least-squares fits of R^2^ vs. time shown in **Figure 3** was 2.00 ± 0.01 (95% confidence interval). The length of the region where Signal was present grew over time because of an increase in Signal concentration down the length of waveguide **(Figure 2b**). Additionally, we observed that at each individual timepoint, R^2^ increased linearly with Carrier concentration, which was predicted by FKPP analysis (supporting information S5).

**Figure 3.**
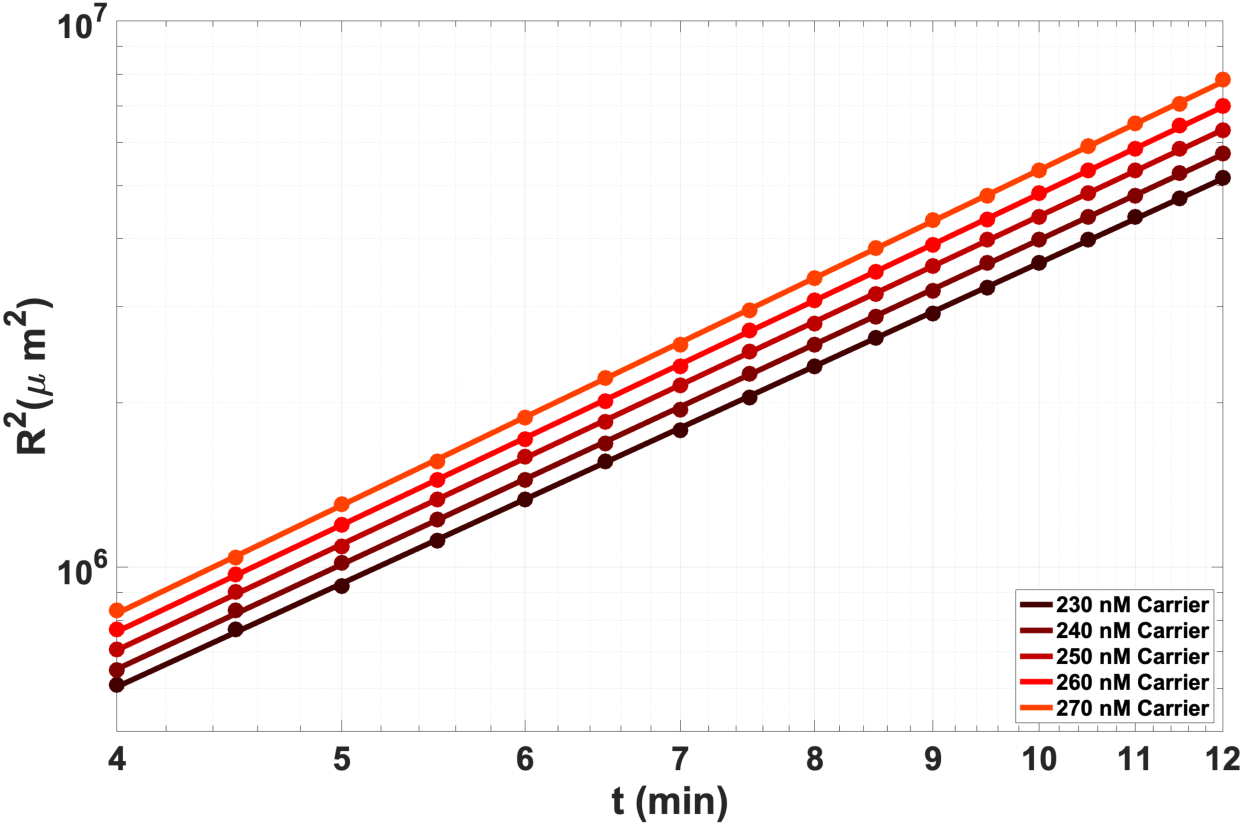
Log scaled plot of the squared wavefront displacement (R^2^) *vs*. time of the idealized waveguide with no Sink present in the core path for varying Carrier concentrations. Dots indicate the squared displacement measured in simulations. Lines are the linear least-squares fit of R^2^ vs. time. A linear relationship exists between log(R^2^) and log(time) at these values.

Having established that the waveguide design could reliably propagate a spatiotemporal wave using known experimental ranges of parameters for DNA strand displacement reactions and diffusion coefficients, we then tested whether it was possible to form a stable traveling wave with Sink patterned within the waveguide core and in the insulation. The inclusion of Sink within the waveguide path provides two key functions. First, Sink can react with Signal at a faster rate than Carrier, so Sink serves as a threshold that can protect the waveguide from spurious activation by leak reactions that produce Signal. Second, the Sink residing within the waveguide removes Signal behind the wavefront, and thus resets the waveguide for future activation.

The simulation used all of the existing conditions described previously and included 35 nM of Sink sequestered within the waveguide core. We observed the formation of a stable traveling pulse in which Signal was degraded into waste at the trailing edge (**Figure 4**). We again observed a nonlinear dependence of R^2^ with time (**Figure 5a**). Logarithmic plots of R^2^ against time yielded a linear relationship across all Carrier concentrations (**Figure 5b**, diamonds & dashed lines). In the presence of 35 nM Sink, the average α calculated by the line of best fit of R^2^ vs. time across all Carrier concentrations and plotted timepoints was 1.71 ± 0.04 (95% confidence interval), indicating that the dynamics of wavefront displacement were still in between the thresholds of directed transport (α = 2) and super-diffusive transport (α > 1). However, the presence of the Sink did significantly impede signal propagation: when 35 nM Sink was in the waveguide core, the R^2^ values at each timepoint were 10-fold smaller than in the absence of Sink.

**Figure 4.**
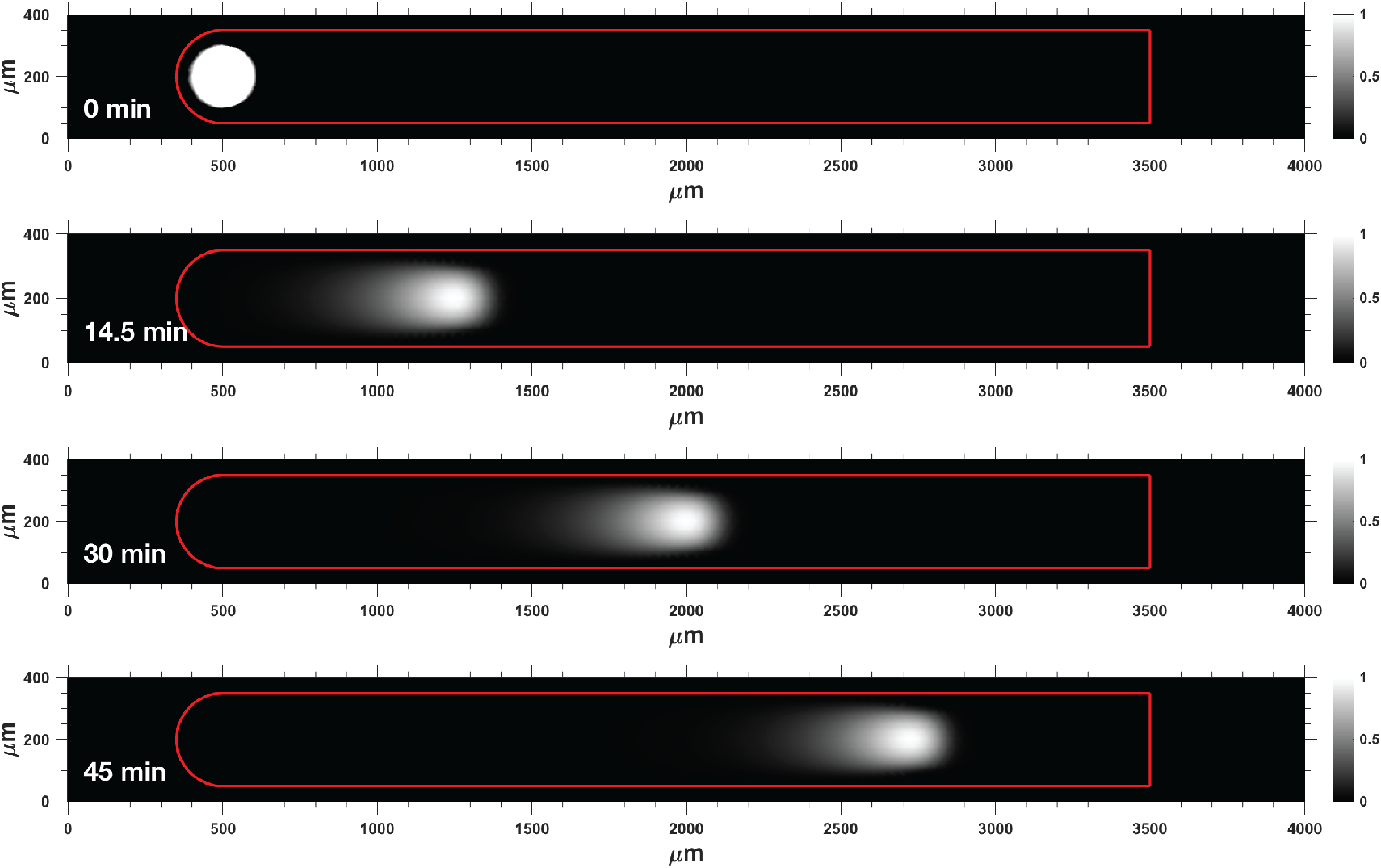
Wave propagation on reaction-diffusion waveguide with 35 nM Sink patterned in the wire core. Spatial propagation of the autocatalytic wavefront over time. Here, Signal trailing the wavefront is eventually degraded. Surface plots are non-dimensionalized by the maximum concentration of Signal within the stable traveling wavefront. The red frame indicates the boundaries of the waveguide.

**Figure 5.**
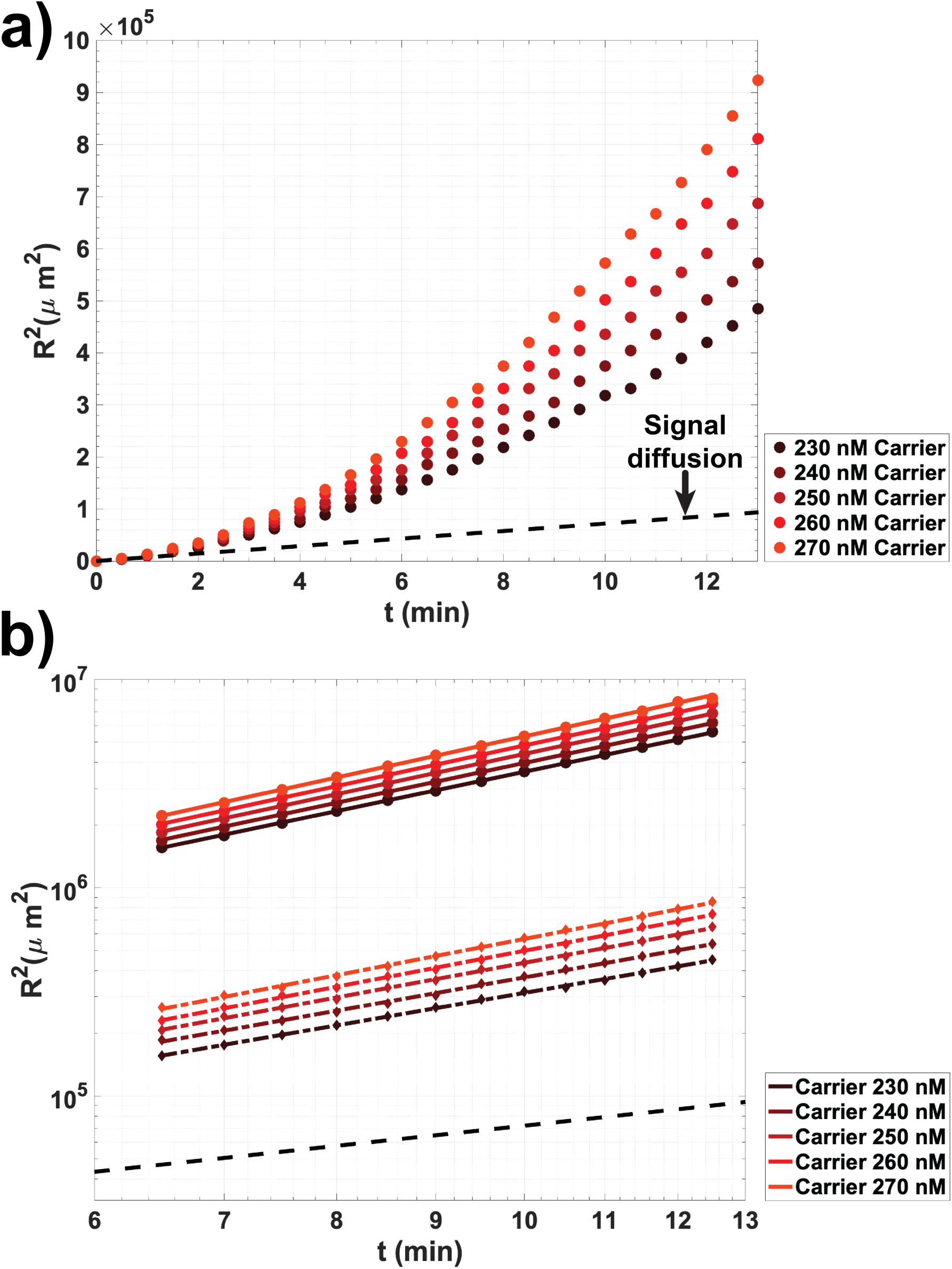
Idealized autocatalytic wavefront propagation in the presence of 35 nM Sink. a) Square of the wavefront displacement, R^2^, vs. time. b) Comparison of R^2^ without Sink (circles are results of PDE reaction-diffusion model & solid lines are the line of best fit for R^2^ vs. time) and with 35 nM Sink (diamonds are the results of the PDE reaction-diffusion model & dashed lines are the line of best fit for R^2^ vs. time) patterned in the waveguide core. Black dashed lines in a) and b) indicate R^2^ for simple diffusion of a 42 nucleotide DNA molecule.

### 3.2 Designing DNA reactions for synthesizing a waveguide: Adding a thresholding reaction to an autocatalytic strand-displacement amplifier mitigates the amplifier’s spurious activation

We next sought to design a chemical reaction network that emulates reactions 1 and 2 using DNA reactions. Designing molecules that react precisely as reactions 1 and 2 was infeasible because the autocatalytic step comprising Reaction 1 cannot be implemented as a single step bimolecular reaction. Reaction 1 was therefore broken into a series of bimolecular strand displacement reactions involving the Carrier species (**Figure 6a**). Minimizing the number and complexity of the reactions required to recapitulate the autocatalytic step provided multiple potential benefits: 1) this limited the number of potential leak reactions that might occur during waveguide operation or when constructing the waveguide, and 2) simplified wire construction with top-down methods such as micro-molding or photolithography. To understand whether DNA strand-displacement reactions could be used to build a DNA waveguide based on these criteria, we modified an autocatalytic DNA strand displacement amplifier previously designed by Zhang and colleagues^44^ to add a thresholding reaction (Figure 6b). We then asked whether the resulting reactions could be used to create a waveguide by executing reaction 1 and 2.

**Figure 6.**
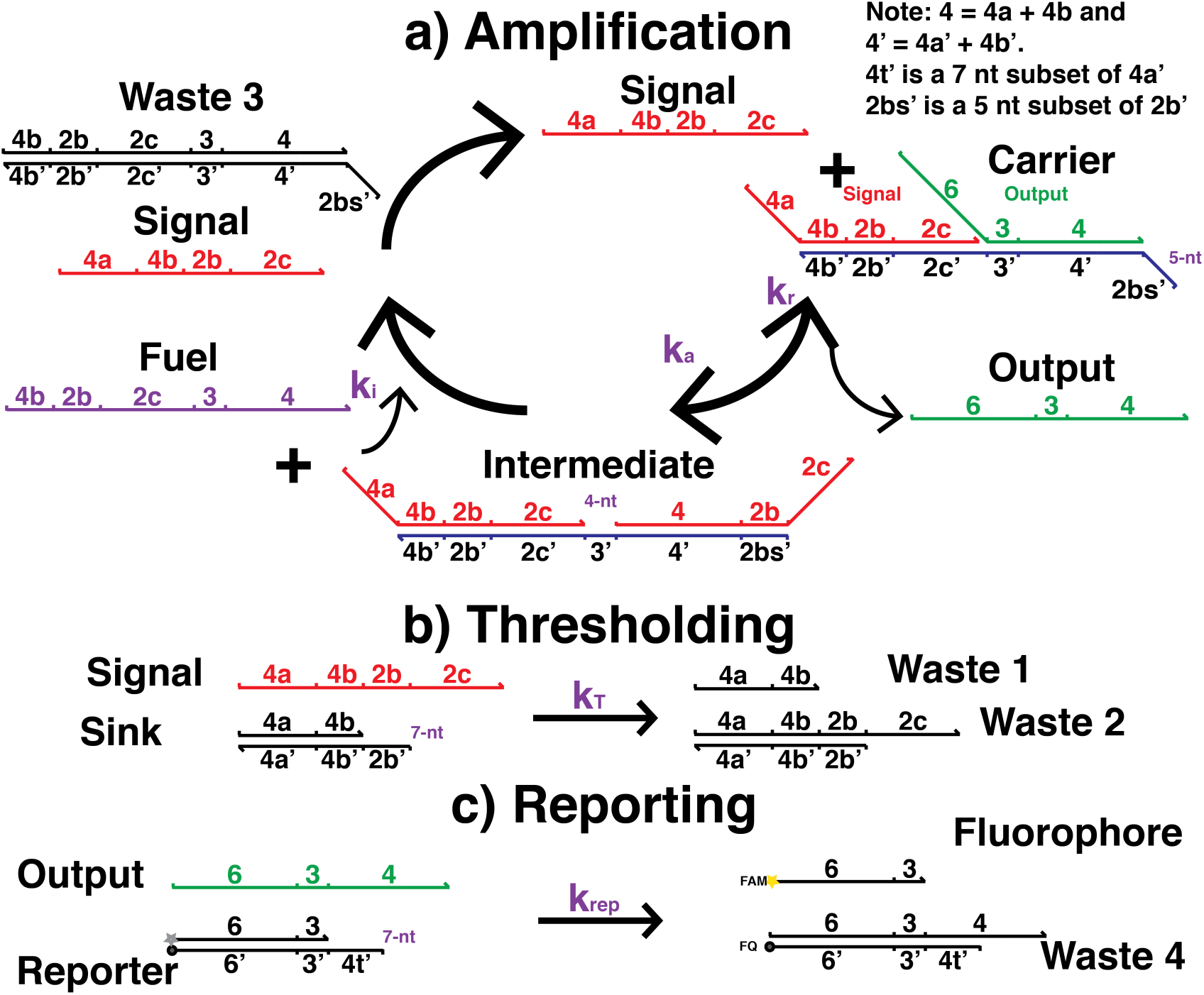
Autocatalytic amplification^44^ reactions with thresholding. a) Autocatalysis module. b) Thresholding reaction module. c) Fluorescence reporting reaction module for optical detection.

The strand displacement implementation of the autocatalytic core of the waveguide consisted of the following reactions: Signal, a single stranded autocatalytic (ss) DNA species, first reacted with Carrier, a complex consisting of Signal and Output strands hybridized to a third strand (**Figure 6a**). After Signal hybridized to the 5 nucleotide (nt) length toehold of this complex, it branch migrated to displace Output, forming Intermediate, a three-strand duplex possessing an exposed toehold (denoted 3’) that Fuel could bind to. The reaction between Signal and Carrier was reversible because Output could also rehybridize to Intermediate and initiate the reverse reaction.

Fuel and Intermediate complex then reacted through a 4-nt toehold and released two Signal strands, each of which then reacted with more Carrier species. Importantly, a large reservoir of Fuel existed within the system, which drove the reaction forward.

To incorporate the insulating and thresholding functions that are key for waveguide function, we created an irreversible thresholding reaction between Signal and a Sink duplex we designed (**Figure 6b**); Signal hybridized to Sink through a 7-nt toehold. We used a Reporting reaction to monitor the progress of the reaction network (**Figure 6c**). The Output strand produced during the amplification cycle reacted with a Reporter duplex composed of a terminal fluorophore-quencher pair to displace its cover Fluorophore strand, enabling optical measurement of the circuit’s reaction progress using quantitative PCR or fluorescence microscopy.

Fuel and Carrier reacted spuriously to produce Signal^42,44^ via base dehybridization at the Carrier duplex terminus and at the nick in the duplex between bound Output and Signal strands. This caused untriggered amplification in the absence of Signal and presented a serious challenge for the use of the amplifier in a spatial system where reactants would be incubated with one another over potentially several hours. We developed a model of the full reaction network in well-mixed conditions to determine the timescale of spurious amplification over a range of concentration conditions. The model was composed of a system of ordinary differential equations (ODEs) and used measured values for the strand displacement reaction rate constants^49,51^ listed in **Figure 6** and for the Carrier-Fuel leak reaction (Supporting Information S3). We observed that for 230-270 nM Carrier incubated with 500 nM Fuel and 50 nM Sink the circuit rapidly entered the growth phase of its sigmoidal activation curve after only 12 minutes (**Figure 7a**). In the absence of any protection chemistry for the Carrier or Fuel species to prevent leakage upon mixing, such a short timescale of activation provided no feasible way for experimental construction of a hydrogel waveguide in a laboratory setting where experimental set up times range from tens of minutes to several hours.

**Figure 7.**
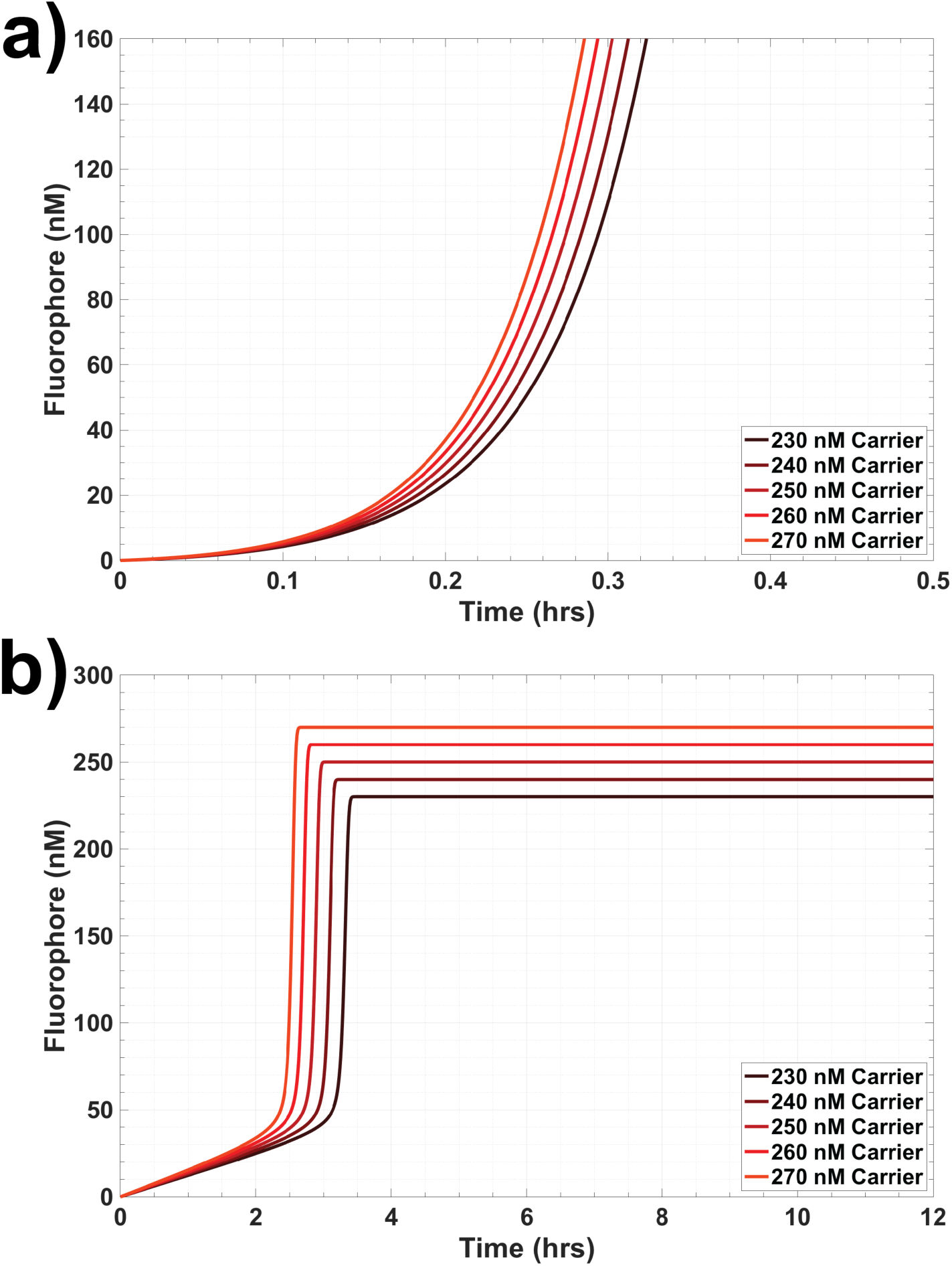
Predictions of a well-mixed reaction model of thresholded autocatalysis. a) The kinetics of amplification when the Carrier’s toehold is 6 nucleotides. b) The kinetics of amplification when the Carrier’s toehold is 5 nucleotides.

To make it possible to build a waveguide, we sought to adapt these reactions in order to increase the time before the autocatalytic reaction proceeded spontaneously (*i*.*e*., the lag time of the circuit). Several strategies were considered to increase the lag phase of the circuit; each entailed performance tradeoffs. Increasing the rate of the threshold reaction by increasing the Sink concentration delays amplification at the cost of raising the threshold concentration of Signal needed to trigger the waveguide. Lowering the rate of amplification by either decreasing the concentrations of Carrier and Fuel or decreasing the rate constant for the amplification reaction would also prolong the lag phase of the reaction at the cost of slowing triggered waveguide operation. We chose to decrease the rate of amplification by shortening the Carrier toehold involved in the reaction between Signal and Carrier from 6 nucleotides to 5 nucleotides, thereby decreasing the rate constant for the reaction by about a factor of 10. With this modification, in the presence of 50 nM Sink, 500 nM Fuel, and varying concentrations of Carrier, the model predicted that the time before the concentration of Signal increased superlinearly was roughly 2.1-2.4 hours (**Figure 7b**). This timescale would allow enough time to build and trigger a waveguide before the leak reaction would become dominant, allowing a proof-of-concept demonstration of the waveguide.

### 3.3 Delayed triggering of autocatalysis

We next sought to test in well-mixed experiments whether it was possible to trigger the circuit by adding a stimulus of Signal while it was held in its lag phase by the Sink reaction. We first measured the time needed for the circuit to reach steady state because of spurious activation, *i*.*e*., when no initial Signal stimulus was added and in the absence of Sink to compare those values with those predicted by our simulations. We mixed a range of concentrations of Carrier (50 nM to 90 nM) with 200 nM Fuel and 150 nM Reporter in the absence of Sink. Reactions were run at 25ºC in a standard reaction buffer (1X Tris-acetate-EDTA buffer with 12.5 mM Mg^2+^). The fluorescence intensity increase of each individual reaction was measured in a Strategene quantitative PCR machine (see supplementary materials, S2). We calibrated and converted fluorescence intensity into Fluorophore concentration using separate calibration wells which were also measured during the experiment (supplementary materials, S4). We defined steady state as the time at which the Flourophore concentration first increased to and remained within 5% of the average concentration attained during the final 15 measurements. Across all Carrier concentrations, we observed that the time to reach steady state was under 40 minutes after the reaction was initiated by the addition of fuel (**Figure 8a**). The steady state times for each reaction condition are listed in **Table 1**. Additionally, the time needed to reach steady state decreased linearly with increasing Carrier concentration. To measure the actual reaction rate constants for the circuit, we fit the rates in an ordinary differential equation model of the amplification circuit to the data using nonlinear least squares regression for each Carrier concentration condition (supplementary materials, S4 and S6 Table S2). The magnitude of the fitted rate constants was consistent with the expected order of magnitude of the rate constants based on toehold length^49^. The expected steady state Fluorophore concentration for each test condition was 50 nM, 60 nM, 70 nM, 80 nM, and 90 nM. The differences between the predictions of the model fit and the measured kinetics are shown in **Figure 8a**. The model was able to fit the measured times to reach steady state as a function of Carrier concentration (**Figure 9a**) to within 8 minutes. Across all conditions, the measured concentration of Fluorophore was slightly greater than the expected steady state concentration predicted by the complete reaction of Fuel and Carrier. The differences in the final Fluorophore concentrations between the model and data are attributed to human pipetting error resulting in variations between calibration and reaction wells.

**Figure 8.**
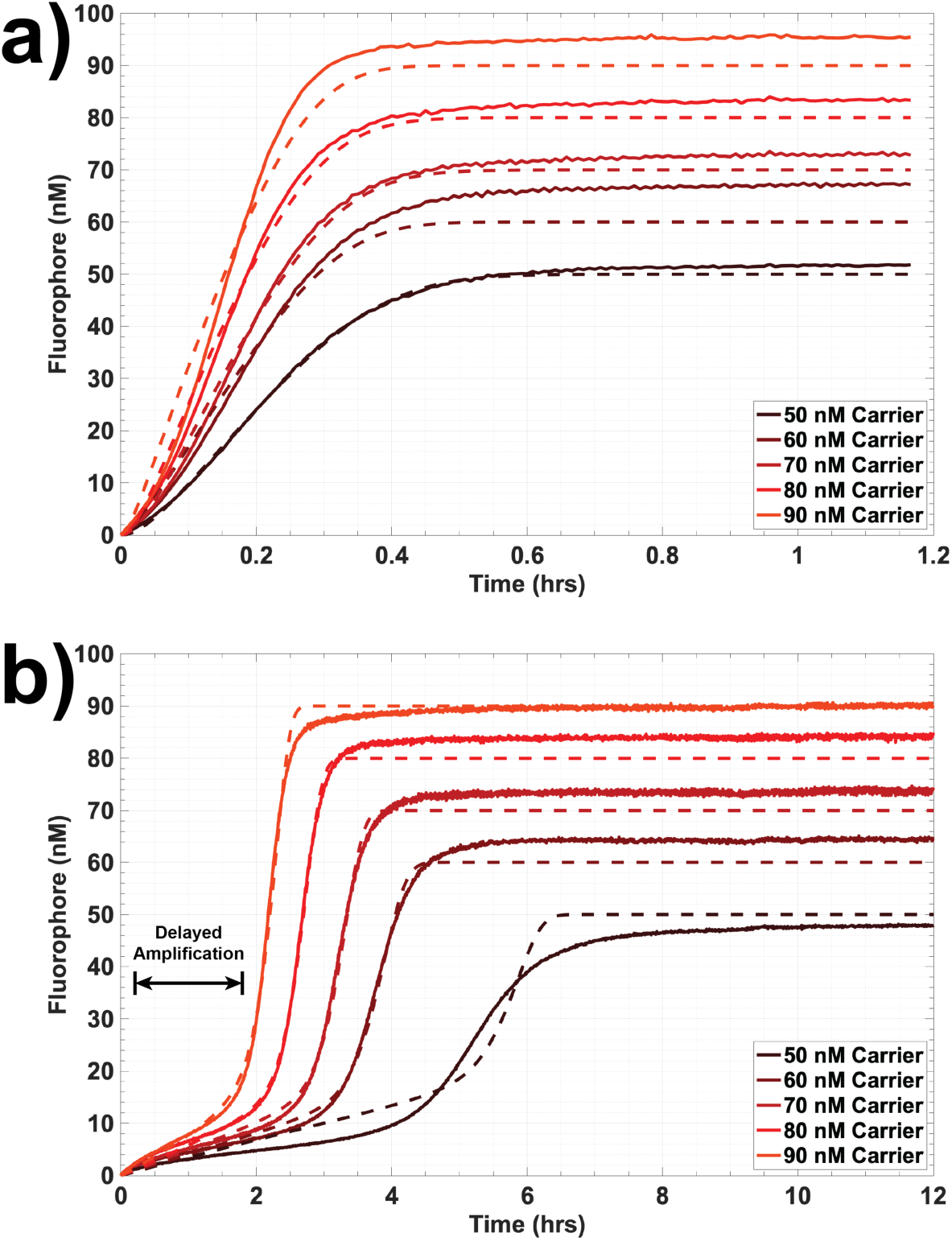
Kinetics of well-mixed autocatalysis reactions in experiments measured by changes in fluorescence. a) Autocatalysis without thresholding by Sink complex and b) autocatalysis in the presence of a thresholding reaction driven by a 50 nM Sink initial condition. Solid lines = experimentally measured concentration profiles, dashed lines = least-squares fit of a reaction model to experimental results.

**Table 1:** Measured Steady State Times for unthresholded and thresholded amplification.

|  | 50 nM Carrier | 60 nM Carrier | 70 nM Carrier | 80 nM Carrier | 90 nM Carrier |
| --- | --- | --- | --- | --- | --- |
| **0 nM Sink** | 30 min | 25 min | 24 min | 21 min | 19 min |
| **50 nM Sink** | 7.3 hrs | 4.7 hrs | 3.9 hrs | 3.2 hrs | 2.7 hrs |
| **X-fold increase** | 15 | 11 | 9.8 | 9.4 | 8.8 |

**Figure 9.**
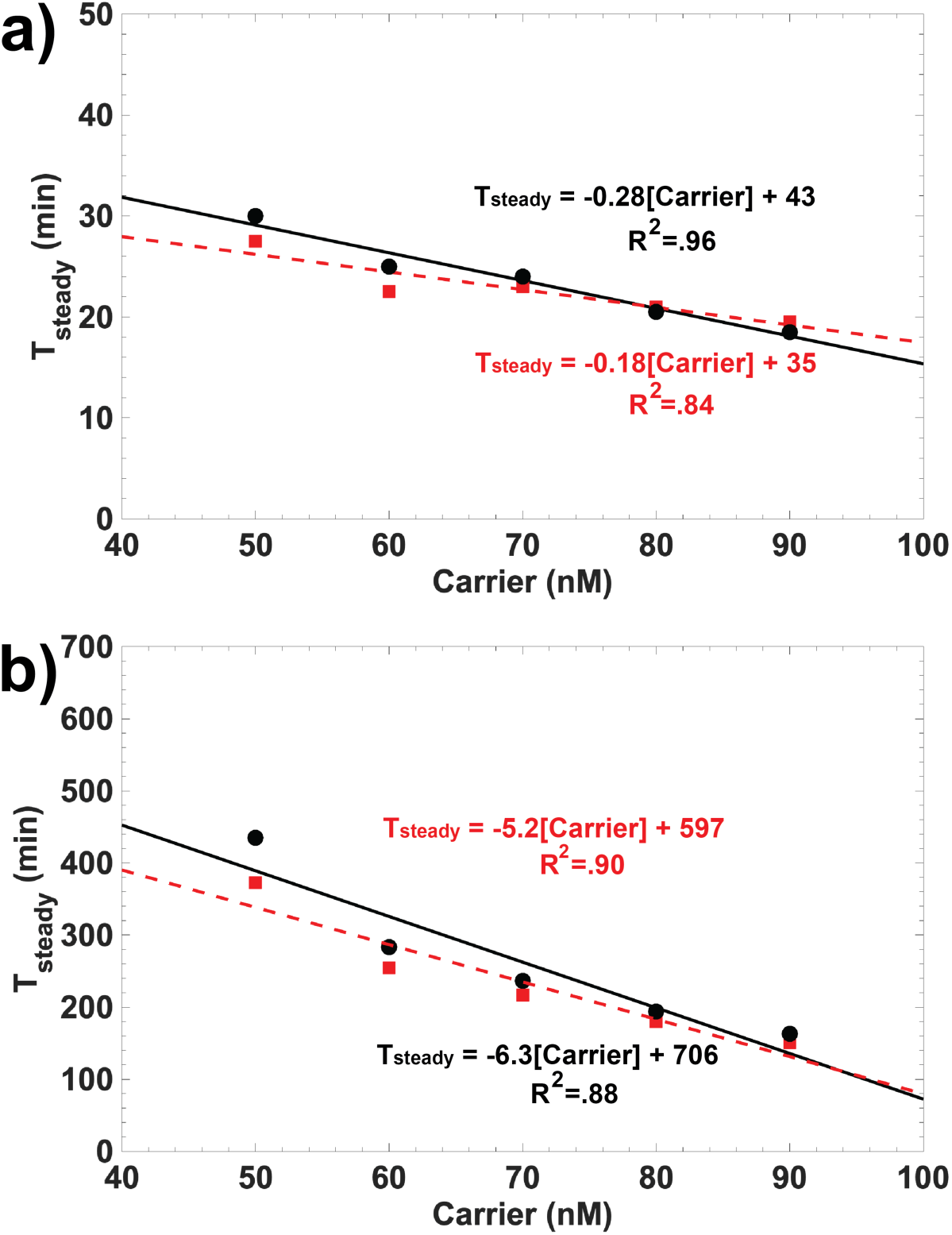
Steady state times for well-mixed autocatalysis: a) without thresholding and b) with 50 nM Sink. Black circles are experimentally measured steady state times. Red squares indicate steady state times predicted by fitting the well-mixed model to experimental data. The solid black line is the linear least squares fit to the experimental steady state times (black circles). The red dashed line is the linear least squares fit to the steady state times predicted by the model (red squares).

Having determined the expected timescale for spurious activation of the unthresholded amplifier, we next tested whether the addition of Sink would delay the onset of amplification and whether a thresholded amplifier could be held in an off state where it could be triggered by an input signal. Importantly, the shape of the Fluorophore curve resulting from thresholded amplification should have a sigmoidal profile. The curve should have an initial period of sub-exponential growth where the threshold quenches autocatalysis by acting as a competitor, followed by autocatalysis after Sink is depleted (**Figure 8b**). Such behavior would indicate that the circuit could eventually undergo exponential growth when a trigger is added, as the trigger would deplete the Sink rapidly. This would then cause the circuit to transition from a lag phase to exponential growth. Conversely, an excess concentration of Sink relative to the Carrier concentration would prevent autocatalysis from occurring and the rate of Output production would only be coupled to the bimolecular reaction of Fuel and Carrier, which would not result in a sigmoidal growth curve. To identify concentrations of Sink and Carrier where autocatalysis would be delayed but not entirely prevented, we repeated the experiments previously described under the same conditions but mixed 50 nM Sink into each reaction well at the start of the experiment. The time needed to reach steady state increased for each reaction with respect to its corresponding process without Sink. The minimum time to reach steady state was 2.7 hours (for the highest Carrier concentration) (**Table 1**). Just as in the absence of Sink, we observed a roughly linear relationship between the time needed to reach steady state and the initial Carrier concentration in the presence of 50 nM Sink (**Figure 9b**). On average across Carrier concentrations, the addition of 50 nM Sink increased the time needed to reach steady state by a factor of 11 ± 3 (95% confidence interval).

We then sought to trigger the circuit during its lag phase by adding Signal to both verify that exponential amplification could occur and identify the size of the Signal stimulus needed to induce exponential amplification. The experimental conditions were identical to those described previously. First, Sink, Fuel, and Reporter were each mixed together in 5 different reaction wells at final concentrations of 50 nM, 200 nM, and 150 nM respectively. Carrier was then added to each of the 5 reaction wells to final concentrations of 50 nM, 60 nM, 70 nM, 80 nM, and 90 nM to initiate the reactions (**Figure 10a**). We then waited 30 minutes, during which time we observed a slow and gradual increase of Fluorophore concentration, consistent with the lag phase of the reaction. After 30 minutes, we added aliquots of Signal corresponding to final concentrations of 20 nM to each of the reactions, we observed sharp increases in the Fluorophore concentration curves, consistent with autocatalytic amplification. These reactions appeared to go to completion: for all Carrier concentrations, the final concentration of Fluorophore was within 7 nM of the concentration expected for complete reaction of Carrier and Fuel. A least-squares fit to the ordinary-differential equation model (**Figure 10a**, dashed lines) was able to match the measured times needed to reach steady state for each of the Carrier concentrations (supporting information, S4). Additionally, the magnitude of the fitted rate constants was consistent with expected the order of magnitude of the rate constants based on toehold length (supporting information, S6 Table S2), suggesting that the bimolecular mechanisms proposed by our model were accurate.

**Figure 10.**
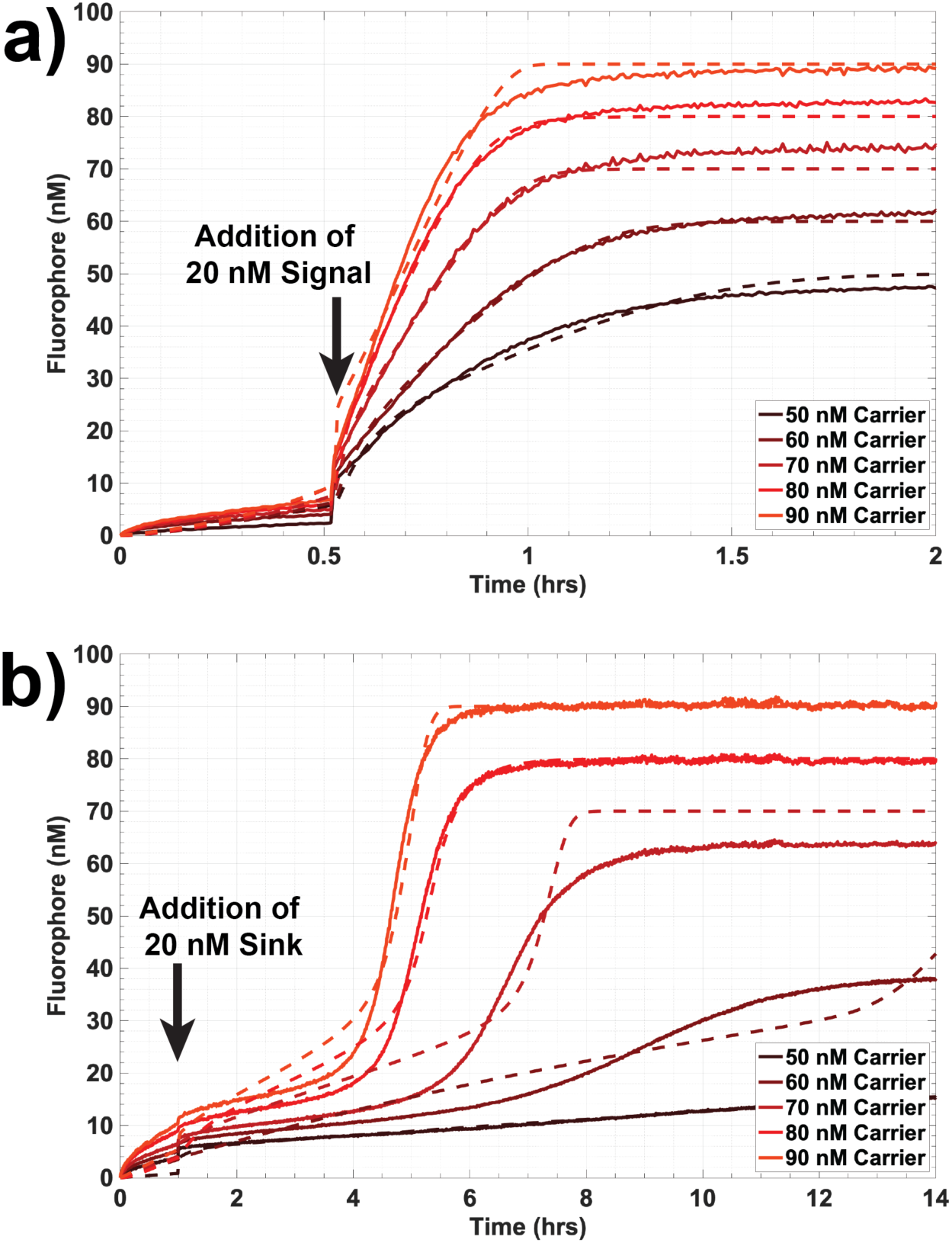
Perturbation of amplification during thresholding. Solid lines = experimental data, dashed lines = results of least-squares regression. a) Delayed triggering of autocatalysis via addition of 20 nM Signal 32 minutes after initiation of the experiment. b) Extended delay of autocatalysis by the addition of 20 nM Sink roughly 1 hr after initiation of the reaction.

To compare this result to the effect of further delaying autocatalysis by adding more Sink, which should provide additional energy to suppress autocatalysis, we conducted the same experiment but added 20 nM of Sink instead of 20 nM Signal 1 hour after initiating the reactions (**Figure 10b**). The addition of 20 nM of Sink increased the total concentration of Sink to 70 nM, which should saturate 50 nM and 60 nM Carrier concentrations and prevent amplification. The fluorescence curves for 60 nM-90 nM Carrier had a sigmoidal shape. At 50 nM Carrier, we observed the slowest increase in Fluorophore across all conditions and no visible inflection of the fluorescence curve, which indicated an absence of exponential growth and inhibition of autocatalysis. This condition did not reach steady state during the timescale of measurement suggesting that the circuit was saturated with Sink. At 60 nM Carrier, we observed a flattened sigmoidal curve indicating that a minimal degree of autocatalysis occurred in the reaction. For this condition, the steady state Fluorophore concentration was below the expected concentration of 60 nM; as a result the least-squares fit of the ODE model overpredicted the timescale of pre-exponential growth after the addition of 20 nM Sink yielding a significant difference between inflection timepoints between the model and experimental curve. Only samples with Carrier concentrations of 80 nM and 90 nM reached their targeted steady state concentrations over the timescale of measurement and had steady state times of 6.1 hours and 5.5 hours respectively, which were both a factor of 2 greater than the steady state times attained by equivalent reactions in the presence of an initial concentration of 50 nM Sink alone. Thus, for 80 nM and 90 nM Carrier, the addition of 70 nM final concentration of Sink mixed into to the circuit at different times before the onset of exponential growth could extend the lag phase.

### 3.4 Mitigation of spurious amplification resulting from leak reactions

Having measured the rate constants of the designed and unintended leak reactions, we then modeled the reaction-diffusion waveguide using the strand displacement reactions in **Figure 6a-b** (see supporting information S3 for CRN implementation). The model initially used the same initial concentration conditions as those stated for the idealized spatial simulations where the Carrier concentration was 270 nM; the initial concentration of Fuel along the waveguide core was 270 nM. The bimolecular rate constants for the strand displacement reaction-diffusion system were based on the expected orders of magnitude which are functions of toehold size.^49^ The results of the initial model are shown in **Figure 11**. In the absence of any Sink within the waveguide core, an initial wave of Signal was observed at 12 minutes. However, the wave decayed at 54 minutes and failed to move forward along the waveguide. Additionally, the spurious generation of Signal from the leak reaction between Fuel and Carrier within the body of the waveguide is evident by 1.5 hrs and grows to turn the whole waveguide on before the wavefront has arrived (**Figure 11a**). When 35 nM of Sink is sequestered within the waveguide core, we observed an initial pulse of Signal at 12 minutes; the wave decayed away by 54 minutes and similarly failed to travel along the waveguide (**Figure 11b**). As a result of these two particular failure modes, we hypothesized that the rate of amplification for the concentration conditions selected was not high enough produce a wave that could overcome the energetic drain of the patterned Sink and insulation.

**Figure 11.**
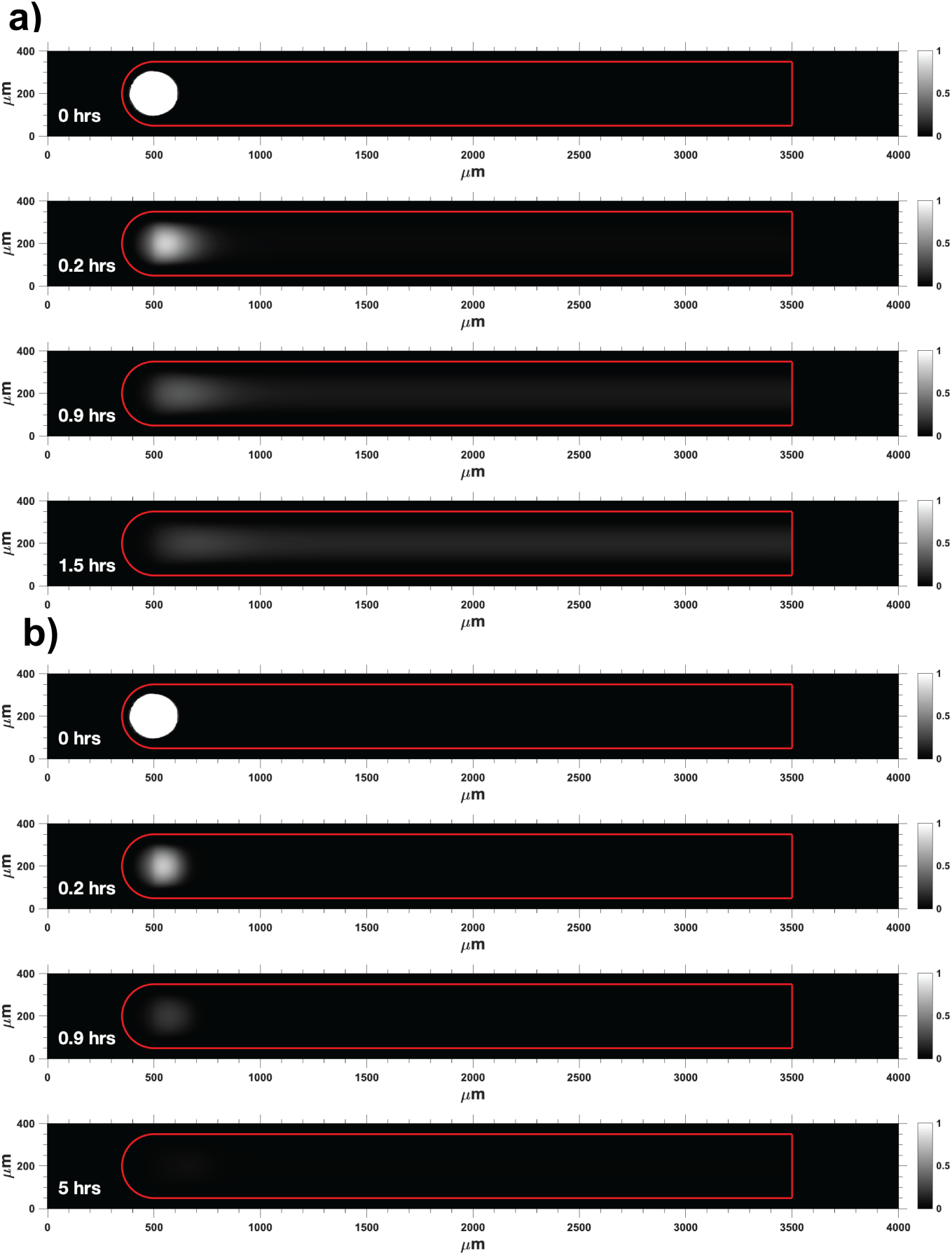
Predictions of a model of spatiotemporal wavefront propagation with non-idealized amplification, thresholding, and Fuel-Carrier leak reactions. a) Propagation failure and spurious waveguide activation without patterned Sink due to Fuel-Carrier leakage. b) Propagation failure with 35 nM Sink patterned within the waveguide core. Surface plots show the Signal concentration divided by the maximum Signal concentration attained on the wavefront.

We next asked whether increasing the rate of amplification would overcome these limitations to enable wave propagation along the waveguide. To achieve this, we modeled a waveguide of the same geometry as the previous simulations. The initial concentrations of Carrier, Fuel and Sink patterned in the waveguide were 650 nM, 650 nM, and 84.24 nM; the ratios of Carrier, Fuel and Sink were kept identical to the earlier simulations of the DNA reaction-diffusion waveguide. The concentration of Sink in the insulation was increased to 1000 nM to prevent the wave from diffusing beyond the waveguide. At 1 hr of simulated time, we observed a wave of Signal that displaced roughly 100 µm from the starting position (Figure 12a). However, at 3 hrs of simulated time, the entire waveguide became active and produced Signal, which corresponded with the complete consumption of Sink, before the wavefront had travelled down the waveguide. Based on this result, we proposed that patterning the reactants as linearly increasing gradients along the length of the waveguide would ensure that Sink was not depleted during the time it took for the wavefront to arrive at a particular location on the waveguide. Additionally, we hypothesized that a linearly increasing gradient would result in a sufficient concentration of Carrier and Fuel at a location ahead of the wavefront so that the wave could continue to propagate and not decay in magnitude or slow its rate of displacement due to the absence of Carrier and Fuel. For example, we modified the simulation so that the concentration of Carrier increased linearly from 780 nM to 1013 nM along the length of the waveguide. The form of the gradients of Fuel, Sink, and Carrier employed in the model are plotted in supporting information Figure S2. The starting concentration of Signal in the Trigger domain was increased to 220 nM and the concentration of Sink in the insulation was increased 1500 nM. We also increased the length of the waveguide to 5000 µm to study the evolution of the wavefront over a longer distance. When the simulation employed linearly increasing gradients of reactants, a wave of Signal propagated along the length of the waveguide (Figure 12b). Analysis of the wave displacement vs. time indicated that the system minimally achieved directed transport, where α = 2.34 ± 0.04 (95% confidence interval) (supporting information Figure S3). These results suggest that patterning Sink, Fuel, and Carrier as linear gradients within the waveguide would serve as an effective strategy for suppressing the autocatalytic leak reaction in single usage experiments.

**Figure 12:**
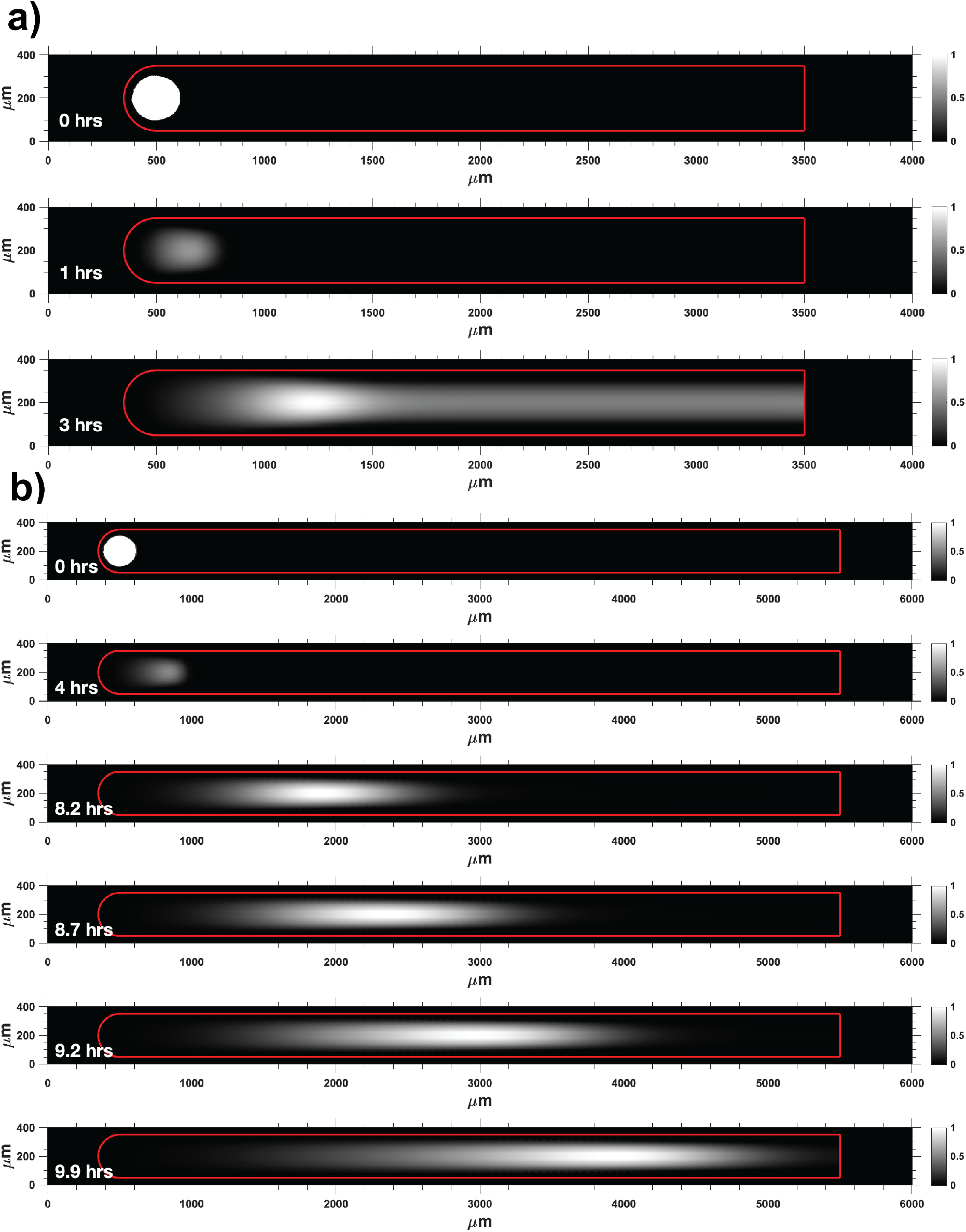
a) Spurious waveguide activation during wavefront displacement due to Carrier-Fuel leak reaction. Initial concentrations of Carrier, Fuel and Sink were 650 nM, 650 nM, 84.24 nM. b) Traveling wave of Signal along waveguide core when Carrier, Fuel and Sink are patterned as linearly increasing gradients.

Moving beyond this analysis, we might ask whether there are molecular protection strategies that might further mitigate the risk of spurious triggering during waveguide construction (*i*.*e*. during photolithographic processes) by keeping the Carrier inactive until its activation is triggered. One protection strategy that could make Carrier inactive to prevent leakage is photo-protection using photocleavable 1-(2-nitrophenyl) ethyl linkers that can be incorporated into the phosphodiester backbone of synthetic oligonucleotides. Such photoprotection of DNA strand displacement reactions using nitro-benzyl chemistries has been demonstrated^52,53^. UV light applied just along the waveguide might be used to photodeprotect Carrier crosslinked to the waveguide just before the waveguide’s use. During testing of the waveguide, we also envision applying this activation just ahead of the Signal as it travels along the length of the wire. Control by photoactivation would minimize the amount of time during which active Carrier and Fuel could react spuriously before the arrival of a wavefront.

We also hypothesized that the dominant mechanism of Carrier-Fuel leak occurred through hybridization of the Fuel strand to transiently exposed nucleotides at the ends of the Carrier duplex, whereby Fuel nucleated to exposed Carrier bases and then branch migrated to displace Signal. In this case, the addition of 7 nucleotide length clamp domains to each end of the Carrier complex would slow the rate of the Carrier-Fuel leak by occluding the ends of the Carrier duplex to prevent invasion by the Fuel. However, these clamping domains would also prevent the desired reaction between Carrier and Signal from occurring during waveguide operation and would require a removal mechanism to yield an active form of the Carrier. UV photocleavable spacers connecting the clamp domains to the 5’ end of Signal and the 3’ end of Output would provide a mechanism for cleaving the clamps to allow autocatalysis to occur at the designed time when the waveguide is exposed to UV light (see Supporting Information: S6 Results & Discussion for photoprotection mechanism).

We experimentally tested whether a clamped Carrier possessing 7 nucleotide length clamp domains would block the leak reaction induced by Fuel (Supporting Information: S6 Results & Discussion for experimental details). However, we observed that incubation of clamped Carrier species with Fuel failed to prevent the leak reaction in well-mixed experiments (supporting information, Figure S5). The persistence of the Carrier-Fuel leak reaction and the size of the fitted leak rate constant indicated that the protection strategy for the duplex ends was not effective in preventing the invasion of Fuel strand. This suggested that the dominant mechanism occurring during the leak reaction was Fuel hybridization to transiently exposed bases at the nick site within Carrier. The proposed hybridization occurred between the 3’end of Signal and the 5’-most nucleotide of Output bound to the bottom strand of Carrier (Carrier_B_); Lysne et al. independently identified the same leak mechanism involving the Carrier nick site^42^. One possible way of occluding the nick to prevent the leak reaction would be to introduce a non-canonical photocleavable attachment between the 5’ end of the last Output nucleotide bound to Carrier_B_ and the 3’carbon of the first Signal nucleotide hybridized to Carrier_B_. However, further study of this mechanism would be needed; to our knowledge, the use of such photosensitive modifications within strand displacement networks has not yet been demonstrated.

## 4 Discussion

Here we use computational analyses and measure the kinetics of reactions in well-mixed solution to support the idea that super-diffusive propagation of chemical waves using DNA strand displacement amplification should be feasible over length scales of hundreds of microns using concentration ranges of oligonucleotides commonly used in strand displacement processes ^54,55^. The use of thresholding reactions provides a way of mitigating deleterious Fuel-Carrier side reactions that might otherwise trigger spurious amplification. The integration of strand displacement waveguides into existing classes of DNA-based soft materials might enable chemical signal transmission within biomaterials and between separated devices over timescales orders of magnitude shorter than what could be achieved with diffusion alone. Moreover, the ability to combine different sets of stimuli using wires will provide control over where and how chemical information is distributed within a biomaterial, enabling coordinated responses to complex sets of environmental cues^56–58^.

To implement a full hydrogel waveguide system experimentally, further investigation of microfabrication methods and nucleic acid photo-chemistries that are DNA-compatible and orthogonal to one another is required. Photolithographic techniques offer the capability of precisely designing patterned biomaterials at biologically relevant size scales within a controlled environment, which is a requirement for strand displacement reactions due to temperature and pH sensitivity. The placement of oligonucleotide patterns within a substrate via photopolymerization to accommodate subsequent photo-directed release or activation of crosslinked species serves as a proxy for spatial biomolecular stimuli that might eventually induce activation of a waveguide and would enable precise spatiotemporal activation of these architectures for further study and optimization.

## Data accessibility

Datasets and MATLAB code supporting this article are available in Dryad Repository through the following link.

## Authors’ contributions

PJD and DS conceived of and designed the study. PJD designed and executed the simulations and experimental work. DS and RS provided technical guidance and analysis. All authors gave final approval for publication and agree to held accountable for the work performed therein.

