## Supplemental Information for "Enabling spatiotemporal regulation within biomaterials using DNA reaction-diffusion waveguides"

### S1. DNA Sequences and Purification

All DNA sequences used in well-mixed experiments were purchased from Integrated DNA Technologies (Coralville, IA).

**Table S1. Waveguide Circuit DNA sequences.**

| Name | Sequence | Purification |
|---|---|---|
| Signal | CATTCAATAC CCTACG TCTCCA ACTAACTTACGG | Desalted |
| Output | ATCCACATACATCATATT CCCT CATTCAATAC CCTACG | Desalted |
| Carrier Bottom | GGAGA CGTAGG GTATTGAATG AGGG CCGTAAGTTAGT TGGAGA CGTAGG | Desalted |
| Sink Cover | CATTCAATAC CCTACG | Desalted |
| Sink Bottom | T TGGAGA CGTAGG GTATTGAATG | Desalted |
| Fuel | CCTACG TCTCCA ACTAACTTACGG CCCT CATTCAATAC CCTACG | Desalted |
| Reporter Bottom | TTGAATG AGGGAATATGATGTATGTGG/3IABKfQ/ | HPLC |
| Reporter Cover | /56FAM/CCACATACATCATATT CCCT | HPLC |
| Clamped Output | CACATAACAA CCACATACATCATATT CCCT CATTCAATAC CCTACG CATACAA | Desalted |
| Clamped Signal | CACCATC CATTCAATAC CCTACG TCTCCA ACTAACTTACGG | Desalted |
| Clamped Carrier Bottom | TTGTATG GGAGA CGTAGG GTATTGAATG AGGG CCGTAAGTTAGT TGGAGA CGTAGG GATGGTG | Desalted |

DNA complexes were annealed in 1X Tris-acetate-EDTA buffer with 12.5 mM $Mg^{2+}$ (TAE/$Mg^{2+}$ buffer). The annealing protocol consisted of heating the solution up to 90 °C for 5 minutes and then cooling 1 °C every minute to 20 °C in an Eppendorf Mastercycler. Annealed complexes were then PAGE (polyacrylamide gel electrophoresis) gel purified to remove single stranded impurities; the conditions were 15% PAGE gels run at 150 V for 3 hours. For the

Carrier complex, two bands were typically observed when visualized at 260 nm; a dark top band was positioned ¼ of the way down the total length of the gel, and a fainter thinner band was located ½ way down the gel length. The top band was cut from the gel and eluted in TAE/$Mg^{2+}$ buffer for 1 day. The eluate was then centrifuged to remove small gel fragments from solution. For the Reporter and Sink complexes, one band was observed during PAGE gel visualization. These bands were cut from the gels, soaked in TAE/$Mg^{2+}$ buffer for 1 day to elute the DNA, and centrifuged to remove small polyacrylamide fragments from solution.

**S2. Well-Mixed Experiments**

All well-mixed kinetic experiments were conducted using a Strategene MX3000 quantitative PCR machine at 25º C. We added reactants to 100 µL total volumes in individual wells of a 96-well plate. The concentrations of reactants listed in the main text are the final concentrations of the species in the 100 µL total volume. Each reaction well contained 1X TAE/$Mg^{2+}$ buffer and 1 µM of PolyT20, a 20 nucleotide poly-thymine strand that acted as sacrificial DNA for adsorption to the polypropylene walls of the reaction wells. To initiate amplification reactions, reactants were added in the following order: Reporter, Carrier, Sink. A baseline fluorescence measurement was then made for 5 minutes. Finally, Fuel and Signal were added to trigger the reaction.

**S3. Modeling of Reaction-Diffusion Waveguides**

Spatial models of reaction-diffusion waveguides were implemented using finite element analysis software using Comsol Multiphysics – Transport of Dilute Species node. The waveguide geometry was meshed with a combination of free tetrahedral and mapped element types. For the idealized waveguide, the model was composed of the following partial-differential equations:

$$\frac{\partial[Signal](t,x)}{\partial t} = D_{ss}\nabla^2[Signal]](t,x) + k_a[Signal](t,x)[Carrier](t,x) - k_d[Signal](t,x)[Sink](t,x)$$

$$\frac{\partial[Carrier](t,x)}{\partial t} = -k_a[Signal](t,x)[Carrier](t,x)$$

$$\frac{\partial[Sink](t,x)}{\partial t} = -k_d[Signal](t,x)[Sink](t,x)$$

Only Signal was allowed to diffuse and it was assigned a diffusion coefficient of 60 μm$^2$ s$^{-1}$, which was the average value measured for a 43 nucleotide sized single stranded oligonucleotide in a 30% (v/v) poly(ethylene-glycol) diacrylate hydrogel[1]. The diffusion coefficients for all other species were set to 0. The full reaction-diffusion waveguide system consists of the following PDEs:

$$\frac{\partial[Signal](t,x)}{\partial t} = D_{ss}\nabla^2[Signal]](t,x) + 2k_i[Fuel](t,x)[Intermediate](t,x) + k_r[Output](t,x)[Intermediate](t,x) - k_a[Signal](t,x)[Carrier](t,x) - k_T[Signal](t,x)[Sink](t,x)$$

$$\frac{\partial[Carrier](t,x)}{\partial t} = -k_a[Signal](t,x)[Carrier](t,x) + k_r[Output](t,x)[Intermediate](t,x) - k_{leak}[Fuel](t,x)[Carrier](t,x)$$

$$\frac{\partial[Sink](t,x)}{\partial t} = -k_T[Signal](t,x)[Sink](t,x)$$

$$\frac{\partial[Output](t,x)}{\partial t} = k_a[Signal](t,x)[Carrier](t,x) - k_r[Output](t,x)[Intermediate](t,x) - k_{rep}[Reporter](t,x)[Output](t,x)$$

$$\frac{\partial[Reporter](t,x)}{\partial t} = -k_{rep}[Reporter](t,x)[Output](t,x)$$

$$\frac{\partial[Intermediate](t,x)}{\partial t} = k_a[Signal](t,x)[Carrier](t,x) - k_i[Fuel](t,x)[Intermediate](t,x) - k_r[Output](t,x)[Intermediate](t,x)$$

$$\frac{\partial[Fuel](t,x)}{\partial t} = -k_{leak}[Fuel](t,x)[Carrier](t,x) - k_i[Fuel](t,x)[Intermediate](t,x)$$

$$\frac{\partial[Fluorophore](t,x)}{\partial t} = k_{rep}[Reporter](t,x)[Output](t,x)$$

The focus of our waveguide analyses was on the autocatalytic species, Signal. To mitigate computational cost and reduce convergence time, the reaction-diffusion network implemented in our Comsol model assumed that Output reacted with an infinitely large source of Reporter throughout the waveguide to instantaneously convert it to Fluorophore. Therefore, the model did not include the reporting reaction of Output and Reporter nor did it incorporate the reverse reaction of Output and Intermediate. Reaction terms not modeled in the Comsol reaction-diffusion simulation are highlighted in purple in the system of PDEs listed above.

**S4. Curve-fitting analysis of data from experiments on the amplifier in well-mixed solution**

Kinetic models of the amplifier were implemented in MATLAB. All fluorescence data were converted from raw fluorescence intensity into Fluorophore concentration by calibrating each experiment. Calibration as performed by adding a known amount of Output to a concentration Reporter within separate individual reaction wells during the experiment. **Figure S2** shows a typical calibration plot. The calibration data allowed us to determine the average proportionality constant, χ, between the average change in fluorescence intensity and the amount of output added:

$$\langle \chi \rangle = \langle \frac{[Output]}{\Delta Counts} \rangle$$

$$[R_f(t)] = \langle \chi \rangle \Delta Counts(t)$$

We then used $\langle \chi \rangle$ to convert all fluorescence counts into Fluorophore concentration. Using this concentration time data, we performed nonlinear least-squares regression using the *lsqcurvefit* Matlab function, which varied reaction rate constants over a range of parameter values to minimize the sum of the square of the y-error between each measured experimental Fluorophore concentration and the Fluorophore concentration predicted by the model. Integration was performed using either the Runge-Kutta method or the variable step variable order method which were implemented using Matlab's *ode45* and *ode15s* functions[2].

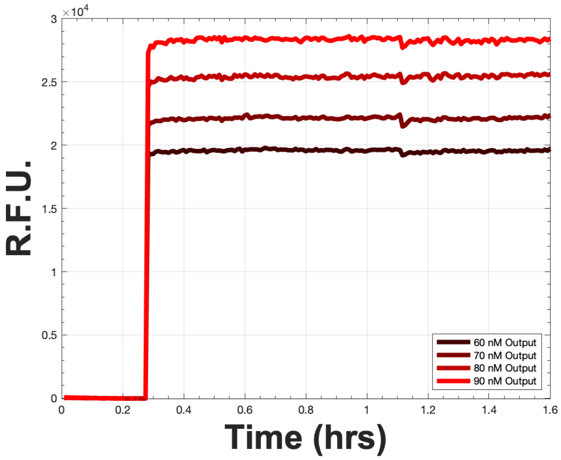


**Figure S1.** An example calibration plot in which different concentrations of Output added to each of 4 reaction wells containing 150 nM Reporter.

These models used the following ODEs describing the reaction rates of the system:

$$\frac{d[Signal](t)}{dt} = 2k_i[Fuel](t)[Intermediate](t) + k_r[Output](t)[Intermediate](t) - [Signal](t)[Carrier](t) - k_T[Signal](t)[Sink](t)$$

$$\frac{d[Carrier](t)}{dt} = -k_a[Signal](t,x)[Carrier](t,x) + k_r[Output](t,x)[Intermediate](t,x) - k_{leak}[Fuel](t,x)[Carrier](t,x)$$

$$\frac{d[Output](t)}{dt} = k_a[Signal](t)[Carrier](t) - k_r[Output](t)[Intermediate](t)$$

$$\frac{d[Reporter](t)}{dt} = -k_{rep}[Reporter](t)[Output](t)$$

$$\frac{d[Sink](t)}{dt} = -k_T[Signal](t)[Sink](t)$$

$$\frac{d[Intermediate](t)}{dt} = k_a[Signal](t)[Carrier](t) - k_r[Output](t)[Intermediate](t) - k_i[Fuel](t)[Intermediate](t)$$

$$\frac{d[Fuel](t)}{dt} = -k_{leak}[Fuel](t)[Carrier](t) - k_i[Fuel](t)[Intermediate](t)$$

$$\frac{d[Fluorophore](t)}{dt} = k_{rep}[Reporter](t)[Output](t)$$

The upper and lower bounds for the fitted rate constants were varied between 4E6 $M^{-1}$ $s^{-1}$ and 0 $M^{-1}$ $s^{-1}$, covering the range of rate constants for bimolecular strand displacement reactions in standard buffer conditions at 25 °C up to a maximum toehold size of 7 nucleotides.

When performing least-squares regression on the amplification perturbation experiments (main text section 3.3, **Figure 10**), our model first integrated the system of ODEs from the starting time to the time of perturbation. At this time point the model took the solution obtained from integration and updated the concentration of Signal or Sink by adding 20 nM of the relevant species to its existing concentration. Numerical integration was continued from the perturbation time to the end of the experiment. The curve fitting function called this model for each specific time point and chose the set of rate constants that minimized the square of the y-error between the model and data set.

**S5 Derivation of Fisher-Kolomogorov-Petrovsky-Piskunov Equation for an autocatalytic wavefront**

For a one-dimensional system, the reaction-diffusion equation describing the accumulation of Signal (abbreviated as Sg below) within the wire over space and time is:

$$\frac{\partial Sg(x,t)}{\partial t} = D_{Sg}\frac{\partial^2 Sg(x,t)}{\partial x^2} + r\big(Sg(x,t)\big) \quad (1)$$

where $D_{Sg}$ is the diffusion coefficient of Signal and r(Sg) is the net reaction rate of Signal, and x is the semi-infinite spatial domain which extends from $x_1$. The initial conditions of the system are:

$$Sg(x,0) = 0 \text{ for all } x < x_1$$

$$Sg(x,0) = Sg_{max} \text{ for all } x \geq x_1$$

The growth rate of Signal is assumed to be bounded:

$$r(Sg_{max}) = 0 \text{ and } r(0) = 0$$

Finally, several restrictions are placed on the growth rate of Signal. First, the reaction rate is assumed to be positive when $0 < Sg(x,t) < Sg_{max}$:

$$r(Sg) > 0$$

Second, the derivative of the reaction rate must satisfy the following inequalities:

$$r'(0) > 0$$

$$r'(Sg) < r'(0) \text{ when } 0 < Sg(x,t) \leq Sg_{max}$$

Far field conditions for the solution to the PDE are:

$$Sg(x,t) \xrightarrow[x \to -\infty]{} 0 \text{ and } Sg(x,t) \xrightarrow[x \to +\infty]{} Sg_{max}$$

We then looked for a solution to the PDE describing an asymptotic traveling wave: $Sg(x,t) = U(z)$, where $z = x + vt$ is a coordinate transformation into one dimension $z$. $z$ reflects the new position of the wave after the passage of time $t$ and rate of displacement $v$. The expression of the reaction-diffusion equation becomes:

$$\frac{v\partial U(z)}{\partial z} = D_{Sg} \frac{\partial^2 U(z)}{\partial z^2} + r\big(U(z)\big) \qquad (2)$$

This second order PDE can then be re-written as a system of first order differential equations. By letting $\frac{dU(z)}{dz} = M$, and substituting M back into equation 2, we get the following expression:

$$M = \frac{dU}{dz} \text{ and } vM = D_{Sg} \frac{dM}{dz} + r(U) \quad (3 \; and \; 4)$$

Equation 4 can be approximated as a linear function of $U$ by recalling that at the unreacted zone immediately preceding the wavefront, the far field condition $U(z) \xrightarrow[z \to -\infty]{} 0$ applies. We can therefore approximate the function $r(U)$ around $U = 0$ by performing a Taylor series expansion of $r(U)$ at this point and inserting the result into eqn. 4:

$$r(U) \approx r(0) + \frac{r'(0)U}{1!} = r'(0)U \quad (5)$$

Equation 4 becomes: $\frac{dM}{dz} = \frac{vM - r'(0)U}{D_{Sg}}$ and the final form of the system of 1st order differential equations becomes:

$$\frac{dM}{dz} = \frac{vM - r'(0)U}{D_{Sg}} \text{ and } M = \frac{dU}{dz} \quad (6 \; and \; 7)$$

This system can also be rewritten back in terms of $U(z)$ as a homogenous constant coefficient 2nd order differential equation:

$$D_{Sg}\frac{d^2U}{dz^2} - v\frac{dU}{dz} + r'(0)U = 0 \quad (8)$$

The exponential solution to this ordinary differential equation will possess the roots of the characteristic equation as exponents. The characteristic equation is:

$$D_{Sg}g^2 - vg + r'(0) = 0 \quad (9)$$

$$g = \frac{v \pm \sqrt{v^2 - 4D_{Sg}r'(0)}}{2D_{Sg}} \quad (10)$$

The roots, $g$, must be real numbers so that the solution of $U(z)$ does not take negative values or exhibit oscillatory behavior. Therefore, the discriminant must be ≥ 0:

$$v^2 - 4D_{Sg}r'(0) \geq 0 \quad (11)$$

By rearranging equation 11, we obtain a requirement for the of the minimum velocity required to from a stable asymptotic traveling wave.

$$v \geq 2\sqrt{D_{Sg}r'(0)} \text{ and } v_{min} = 2\sqrt{D_{Sg}r'(0)} \quad (12 \text{ and } 13)$$

It is important to note that the minimum rate of displacement does not depend on the initial conditions of the system. Additionally, $r'(0)$ can be determined for the for the autocatalytic circuit discussed previously in the absence of Sink:

$$r'(U(z)) = r'(Sg(x,t)) = \frac{\partial}{\partial Sg}[k_aC \times Sg] = k_a\left(C\frac{\partial Sg}{\partial Sg} + Sg\frac{\partial C}{\partial Sg}\right) \quad (14)$$

$$r'(0) = k_aC_{max}$$

$$v \geq 2\sqrt{D_{Sg}k_aC_{max}} \text{ and } v_{min} = 2\sqrt{D_{Sg}k_aC_{max}} \quad (15 \text{ and } 16)$$

where the net reaction rate of Signal is differentiated with respect to Signal using the product rule and evaluated at [Signal] = 0; note that we assumed that at the leading edge of the wavefront where Signal approaches 0, Carrier takes its maximum concentration value, $C_{max}$. In the presence of Sink (Sk), $r'(0) = k_aC_{max} - k_dSk_{max}$. This leads to the expressions:

$$v \geq 2\sqrt{D_{Sg}(k_a C_{max} - k_d S k_{max})} \text{ and } v_{min} = 2\sqrt{D_{Sg}(k_a C_{max} - k_d S k_{max})} \quad (17 \textit{ and } 18)$$

### S6. Results & Discussion

**Molar free energy change during strand displacement amplification:**

The total Gibbs free energy change of the reaction can be expressed as the sum of the standard free energies of the species produced minus sum of the standard free energies of species consumed:

$$\Delta G_{rxn} = \sum_i y_i \, \Delta G^{\circ}_{product\ i} - \sum_i x_i \, \Delta G^{\circ}_{reactant\ i}$$

where $\Delta G^{\circ}_i$ is the molar free energy of a particular DNA species and $y_i$ and $x_i$ are the number of moles produced or consumed during the reaction step. The total reaction for 1 cycle of amplification is:

$$Carrier + Fuel \xrightarrow{k} Signal + Output + Waste_3 \; (SI\ Rxn. S. 1)$$

The molar Gibbs free energy for each species at 25 ºC in standard buffer conditions can be calculated using the nearest-neighbor model for DNA structural motifs[3], which assumes that the energy of the species is determined by the composition and locations of its base pairs. For DNA duplexes, each base-pair within the duplex is assigned a standard free energy based on the base pairing interaction (A-T/G-C), and the base-pairs directly adjacent to it to account for base stacking interactions. Additional factors for duplex stability accounted for by the model are the presence of terminal A-T and G-C pairings, the entropic penalty associated with nucleation of the first base-pair, and coordination of counter-ions with the backbone, which are all accounted for together with an initiation/terminal base-pairing term, and a symmetry term if the duplex is self-complementary. Together the standard free energy of each species can be expressed as:

$$\Delta G^{\circ}_i = \sum_j n_j \Delta G^{\circ}_j + \Delta G^{\circ}(init. term\ G-C) + \Delta G^{\circ}(init. term\ A-T) + \Delta G^{\circ}_{sym}$$

$\Delta G_j^\circ$ is the standard free energy for the $n_j$ possible base-pairs in the species. The values for these free energies have been computed and correlated across a variety of temperature and salt conditions[4–6]. Here, we used software tools, specifically NUPACK[7] to calculate the free energy of each species at standard reaction conditions; the NUPACK assumptions were 25 ºC in a buffer containing 12.5 mM $Mg^{2+}$ and 1 M $Na^+$.

$$\Delta G^{\circ}_{Carrier} = -72.43\ kcal\ mol^{-1}$$

$$\Delta G^{\circ}_{Waste_3} = -73.10\ kcal\ mol^{-1}$$

$$\Delta G^{\circ}_{Signal} = -2.21\ kcal\ mol^{-1}$$

$$\Delta G^{\circ}_{Output} = 0.0\ kcal\ mol^{-1}$$

$$\Delta G^{\circ}_{Fuel} = -2.21\ kcal\ mol^{-1}$$

$y_i$ and $x_i$ = 1 for all species in SI reaction S1. Therefore, we expect $\Delta G_{rxn} = -0.67\ kcal\ mol^{-1}$ for the completion of 1 cycle of amplification at 25º C in the presence of 12.5 mM $Mg^{2+}$. For comparison, the average molar thermal energy fluctuation from molecular collisions at 25 ºC is $kT * N_A = 0.59\ kcal\ mol^{-1}$, where $k$ is the Boltzmann constant and $N_A$ is Avogadro's number, illustrating how close the free energy change of the system is to the energy provided by random molecular collisions.

**Measurement of Amplifier Rate Constants**

We first fit reaction rate constants in well-mixed conditions for the un-thresholded amplifier. The fitted parameters were the reaction rate constants $k_a$, $k_r$, $k_{rep}$ and $k_i$ shown in the main text reaction diagram Figure 6. The strand displacement mechanism for the reaction of Fuel and Intermediate and Signal and Intermediate occur through the same toehold and involve branch migration along specificity domains of roughly equal length, so we assumed that the rate constants $k_r$ and $k_i$ were equal in our model. The average values for the fitted parameters are listed in **Table S2** and the least-squares fit for each reaction is plotted as a dashed line in main text Figure 8. We observed that the estimated magnitudes of $k_{rep}$, $k_r$ and $k_i$ from the model were within an order of magnitude of known experimental ranges for reactions with the corresponding toehold sizes. The expected magnitudes of 7 nucleotide, 6 nucleotide, and 4 nucleotide toehold bimolecular rate constants are 3E6 $M^{-1}$ $s^{-1}$, 5E5 $M^{-1}$ $s^{-1}$, 5E3 $M^{-1}$ $s^{-1}$ respectively[8]. Interestingly, the magnitude of fitted rate constant $k_a$ (which involved a 5 base-pair toehold ~ $10^4$ $M^{-1}$ $s^{-1}$) was higher than the expected value by a factor of 10. Additionally, the measured leak rate constant for the leak reaction between Fuel and Carrier was ~ $10^3$ $M^{-1}$ $s^{-1}$, roughly 2 orders of magnitude higher than the value previously reported by Zhang et al.[9]. Key differences exist between the purity of the strands used in their experiments and in our experiments. Zhang et al. used HPLC purified DNA. All non-modified strands purchased from IDT in our experiments were ordered with standard desalting, which can yield a higher fraction of oligonucleotides with 5'end nucleotide deletion errors than what is found in HPLC purified DNA. 5' deletion errors could expose bases at the end of the 4b' domain of Carrier, effectively creating a permanent 1 or 2 nucleotide toehold for Fuel to hybridize to, in addition to the Carrier nick, and opposite duplex end which both offer possible invasion points for Fuel. These toeholds could account for the higher observed reaction rates. Finally, subtle differences also existed between the duplex purification protocols used in both experiments. Zhang et al. purified DNA duplexes using 12% non-denaturing polyacrylamide gel electrophoresis gels using a power of 180V for 6 hours. Our protocol used 15% non-denaturing polyacrylamide gel electrophoresis gels run at 150V for 3 hours.

Similarly, the average rate constants fitted to the thresholded amplifier data yielded a similar trend to what was observed with the unthresholded system. Here, we fit $k_a$, $k_r$, $k_{rep}$, $k_i$,

$k_{leak}$, and $k_t$. We observed that the magnitudes of $k_r$ and $k_i$ were in the expected range for a 4-nt toehold reaction. However, the predicted magnitude of $k_a$ was an order of magnitude higher than its expected value. Additionally, $k_{rep}$ was one order of magnitude lower than the expected rate constant for a reaction involving a 7 nucleotide toehold rate constant ~ $10^6$ $M^{-1}$ $s^{-1}$ while $k_t$ was on the expected order of a 7 nucleotide rate constant. Finally, the magnitude of $k_{leak}$, which was 85 $M^{-1}$ $s^{-1}$, fell within the expected range for a reaction involving a 0-2 nucleotide toehold ~ 10-100 $M^{-1}$ $s^{-1}$. It is important to note that during purification of the Carrier complex, it was incredibly difficult to ensure consistency in the fraction of properly formed complex; different experiments used different batches of purified Carrier. Variation between these results across data sets may be attributed to differences in Carrier purity from batch to batch as was observed by Zhang et al.[9].

**Table S2: Un-thresholded and thresholded amplifier average fitted rate constants (95% confidence intervals)**

| | **ka** | **kr** | **ki** | **kt** | **krep** | **kleak** |
|---|---|---|---|---|---|---|
| **0 nM Sink** | 1.9E5 ± 1.5E4 $M^{-1}$ $s^{-1}$ | 8.9E3 ± 1.6E2 $M^{-1}$ $s^{-1}$ | 8.9E3 ± 1.6E2 $M^{-1}$ s-$^{1}$ | N/A | 1.8E5 ± 1.5E5 $M^{-1}$ $s^{-1}$ | 2.9E3 ± 5.0E2 $M^{-1}$ $s^{-1}$ |
| **50 nM Sink** | 1.2E5 ± 3.2E4 $M^{-1}$ $s^{-1}$ | 6.8E3 ± 1.8E3 $M^{-1}$ $s^{-1}$ | 6.8E3 ± 1.8E3 $M^{-1}$ $s^{-1}$ | 3.8E6 ± 2.4E5 $M^{-1}$ $s^{-1}$ | 9.7E5 ± 1.1E5 $M^{-1}$ $s^{-1}$ | 8.5E1 ± 1.4E0 $M^{-1}$ $s^{-1}$ |

**Table S3: Average fitted rate constants for thresholded amplifier perturbation experiments (95% confidence intervals).**

| | **ka** | **kr** | **ki** | **kt** | **krep** | **kleak** |
|---|---|---|---|---|---|---|
| **20 nM Signal Addition** | 1.9E6 ± 7.7E5 $M^{-1}$ $s^{-1}$ | 5.2E3 ± 3.1E3 $M^{-1}$ $s^{-1}$ | 5.2E3 ± 3.1E3 $M^{-1}$ $s^{-1}$ | 7.7E5 ± 4.8E5 $M^{-1}$ $s^{-1}$ | 2.6E5 ± 5.9E5 $M^{-1}$ $s^{-1}$ | 1.0E2 ± 5.4E1 $M^{-1}$ $s^{-1}$ |
| **20 nM Sink Addition** | 1.1E5 ± 8.1E4 $M^{-1}$ $s^{-1}$ | 7.9E3 ± 1.4E3 $M^{-1}$ $s^{-1}$ | 7.9E3 ± 1.4E3 $M^{-1}$ $s^{-1}$ | 1.6E6 ± 8.2E5 $M^{-1}$ $s^{-1}$ | 1.3E6 ± 2.0E6 $M^{-1}$ $s^{-1}$ | 6.7E1 ± 3.3E1 $M^{-1}$ $s^{-1}$ |

Fitted rate constants for the perturbation experiments are listed in **Table S3**. We again observed that the optimized magnitudes for the rate constants corresponded to toehold sizes that were within 1 nucleotide of with the actual sizes involved in the experimental system.

**Assessment of a Photoprotection Strategy for prevention of Carrier-Fuel leak reactions**

As a first attempt to mitigate the leak reaction between Carrier and Fuel in the amplification reaction, we chose to add 7-bp clamp domains to both ends of the duplex that had no complementarity to Fuel. To prevent Fuel from reacting with Carrier, we extended the length of Signal and Output to contain the reverse complement of the 7 nucleotide domains added to the bottom strand of Carrier (referred to as $Carrier_B$). Importantly, the original unclamped sequence structure of Signal and Output, and $Carrier_B$ was retained. The toehold of $Carrier_B$ and 4a domain of Signal formed bulge loops (**Figure S2**) in the duplex. The hypothesis of this design was that: 1) the presence of clamps would slow the rate at which Fuel could nucleate with frayed bases at the ends of $Carrier_B$ due to steric hindrance, and that 2) during partial displacement of Signal or Output by Fuel, the clamps would increase the rate of rehybridization and reverse branch migration of Signal and Output because these molecules possess a domain to reattach and/or remain attached to Carrier duplex, thereby forcing these oligos into a set of conformational configurations that lower the energy barrier for base nucleation with adjacent segments of Fuel-hybridized duplex. During the photo-deprotection process, Signal and Output would be attached to their clamp domains with 1-(2-nitrophenyl) ethyl linkers (**Figure S2**). Exposure of Carrier to UV light would break these linkages and produce the functional form of Carrier where Signal and Output can be fully displaced from the complex during strand displacement. To verify that the protected form of Carrier, $Carrier_p$, reacted with Fuel at a slower rate or did not react at all, we first mixed 50 to 90 nM $Carrier_p$ with 200 nM Fuel and 150 nM Reporter in different reaction wells of a 96 well plate. We tracked the increase in Fluorophore concentration over time (**Figure S3**) and observed a slow and gradual increase in Fluorophore concentration, where the rate of increase over time appeared to be proportional to the initial Carrier concentration. Additionally, all kinetic traces maintained their concavity and no inflection points were visually observed over the timescale of measurement, suggesting that autocatalysis was inhibited and the rate of Fluorophore production was largely coupled to the bimolecular reaction of Fuel and $\text{Carrier}_p$. Based on these observations, we designed a ODE model of the reaction which assumed that autocatalysis was inhibited (i.e. Signal could not react with $\text{Carrier}_p$) and that Fuel was able to react with $\text{Carrier}_p$ to produce Output (see Supporting Information: Materials & Methods for model equations). Nonlinear least-squares regression was performed to fit the model to the experimental data: $k_{leak}$ and $k_{rep}$ were the fitted parameters. The average values of $k_{leak}$ and $k_{rep}$ were 22 ± 1.9 $M^{-1}$ $s^{-1}$ and 2.8E6 ± 1.5E6 $M^{-1}$ $s^{-1}$, which were within one order of magnitude of values obtained from previous fitting analyses of the leak and reporting reaction rates.

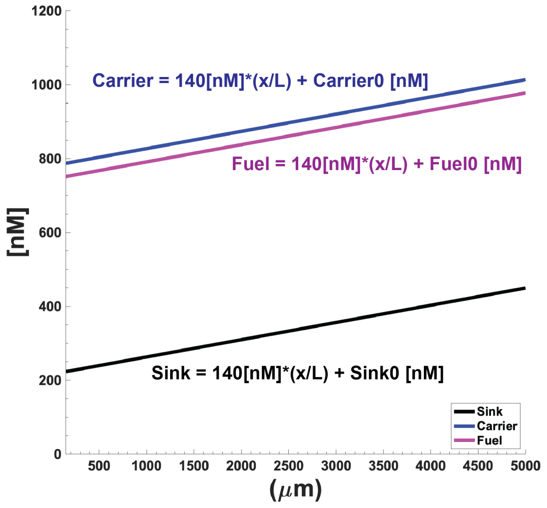


**Figure S2.** Initial concentration gradients of Carrier, Fuel, and Sink employed in the gradient patterned DNA reaction-diffusion waveguide.

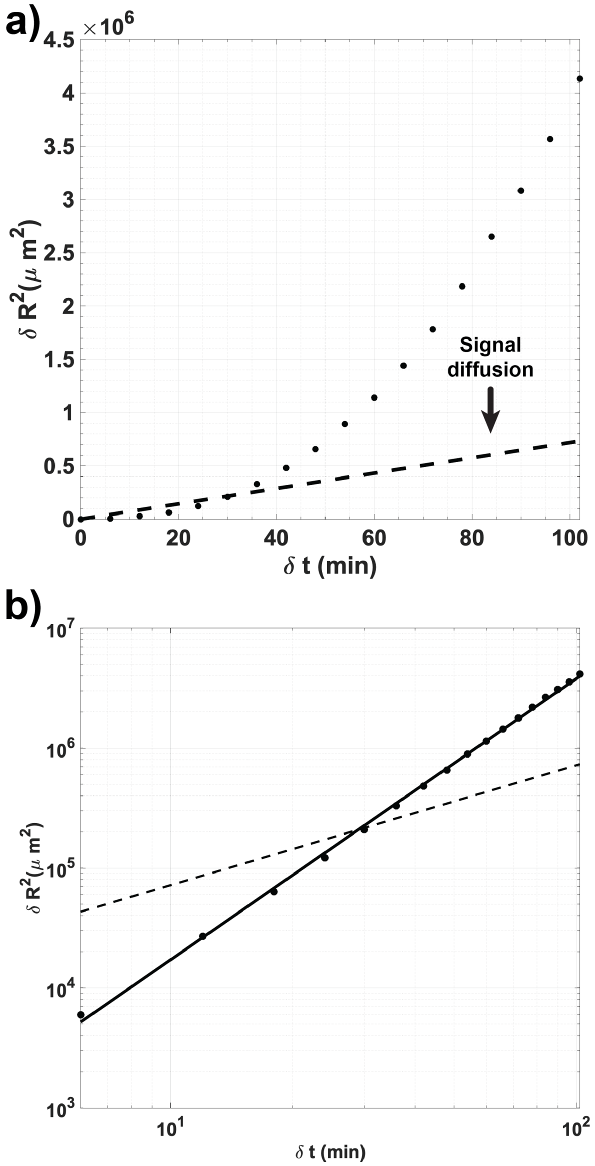


**Figure S3.** Relative wave displacement vs. time. a) Linear scale. b) Log-log scale. Black circles are results of PDE reaction-diffusion model and the solid black line represents the line-of-best-fit. Average slope = 2.34 ± 0.04 (95% CI). Black dashed lines in a) and b) indicate the expected mean-squared displacement for diffusion of a 42 nucleotide DNA molecule. The simulated time window analyzed here is 8.2 hrs - 9.9 hrs in the Comsol model.

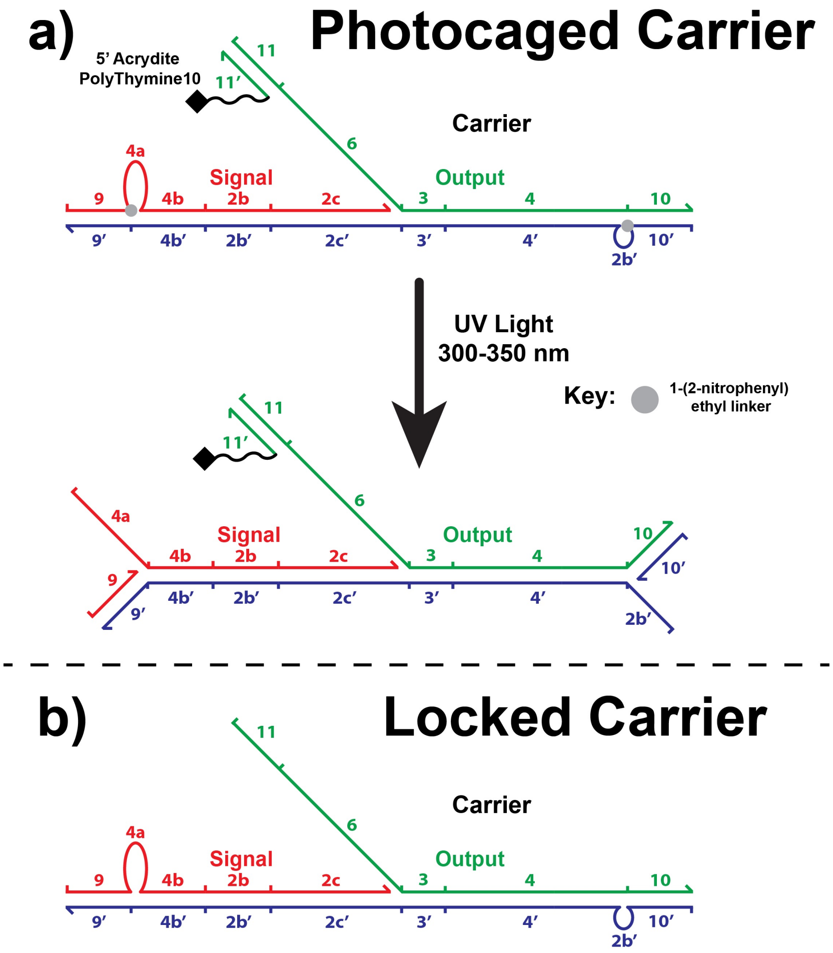


**Figure S4.** a) Carrier photoprotection strategy using nitrobenzyl-modified clamp domains to prevent Fuel leakage with Carrier duplex ends. Photocleavage of 1-(2-nitrophenyl) ethyl linkers results in exposure of the 2b' toehold on Carrier and the activation of bound Signal. b) Locked Carrier substrate tested in well-mixed experiments for its ability to slow the Fuel-Carrier leak reaction.

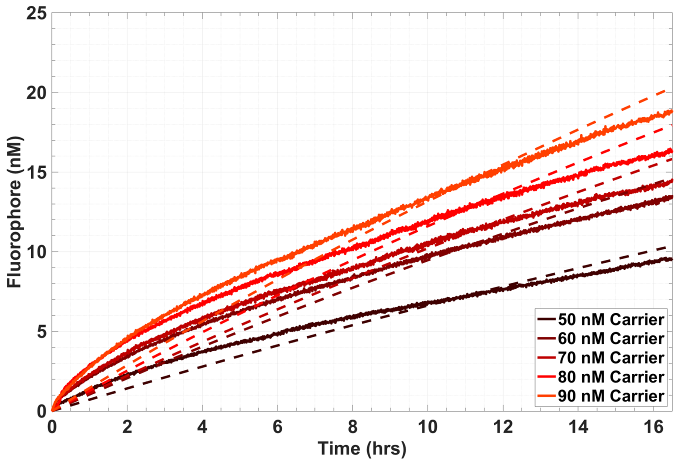


**Figure S5.** Fluorescence signals generated from incubation of 50-90 nM Locked Carrier with 200 nM Fuel.

(8) Zhang, D. Y.; Winfree, E. Control of DNA Strand Displacement Kinetics Using Toehold

Exchange. *J. Am. Chem. Soc.* **2009**, *131*, 17303–17314.

(9) Zhang, D. Y.; Turberfield, A. J.; Yurke, B.; Winfree, E. Engineering Entropy-Driven Reactions and Networks Catalyzed by DNA. *Science (80-. ).* **2007**, *318* (5853), 1121–1125.
